# Structure of intact human MCU supercomplex with the auxiliary MICU subunits

**DOI:** 10.1101/2020.04.04.025205

**Authors:** Wei Zhuo, Heng Zhou, Runyu Guo, Jingbo Yi, Lei Yu, Yinqiang Sui, Laixing Zhang, Wenwen Zeng, Peiyi Wang, Maojun Yang

**Author notes:** These authors contribute equally to this work (Wei Zhuo, Heng Zhou, Runyu Guo, Jingbo Yi). This Paper is submitted back-to-back with the following paper:* V. Garg et al., The Mechanism of MICU-Dependent Gating of the Mitochondrial Ca2+ Uniport er. bioRxiv 2020.04.04.025833; doi: https://doi.org/10.1101/2020.04.04.025833.

## Abstract

The mitochondrial Ca^2+^ uniporter (MCU) supercomplex is essential for mitochondrial Ca^2+^ uptake. Here, we present high-resolution cryo-EM structures of human MCU-EMRE supercomplex (MES, 3.41 Å) and MCU-EMRE-MICU1-MICU2 supercomplex (MEMMS, 3.64 Å). MES adopts a V-shaped dimer architecture comprising two hetero-octamers, and a pair of MICU1-MICU2 hetero-dimers form a bridge across the two halves of MES to constitute an O-shaped architecture of MEMMS. The MES and MEMMS pore profiles are almost identical, with Ca^2+^ in the selectivity filters and no obstructions, indicating both channels are conductive. Contrary to the current model in which MICUs block the MCU pore, MICU1-MICU2 dimers are located on the periphery of the MCU pores and do not occlude them. However, MICU1-MICU2 dimers may modulate MCU gating by affecting the matrix gate through the EMRE lever.

## Introduction

Mitochondrial Ca^2+^ homeostasis regulates energy production, cell division, and cell death. The basic properties of mitochondrial Ca^2+^ uptake have been firmly established (*1-4*). The Ca^2+^ influx is mediated by MCU, driven by membrane potential and using a uniporter mechanism (Ca^2+^ transport is not coupled to transport of any other ion) (*5*). Patch-clamp analysis of MCU currents demonstrated that MCU is a channel with exceptionally high Ca^2+^ selectivity (*6*). Further, Ca^2+^ efflux is known to involve two pathways: H^+^/Ca^2+^ exchange (*7*) and Na^+^/Ca^2+^ exchange (*8*). The homeostasis of mitochondrial Ca^2+^ must be exquisitely regulated to prevent wastage of energy from bidirectional Ca^2+^ flux (*9-11*).

In 2010, an RNA silencing study highlighted that MICU1 (mitochondrial Ca^2+^ uptake 1) represents the founding member of a set of proteins required for high-capacity mitochondrial Ca^2+^ uptake (*12*). In 2011, two groups independently reported the membrane protein MCU and proposed it as the pore-forming element of the long-sought mitochondrial Ca^2+^ uniporter (*13, 14*), which was confirmed later using whole-mitoplast voltage-clamping (*15*). In 2012, a paralog of MICU1, MICU2 (mitochondrial Ca^2+^ uptake 2), was shown to cooperatively regulate Ca^2+^ uptake with MICU1 (*16*). In 2013, a genetic study led to the characterization of MCUb (mitochondrial Ca^2+^ uniporter b), a vertebrate specific protein sharing ∼ 50% sequence identity and the same membrane topology with MCU (*17*). Also in 2013, a previously uncharacterized protein, EMRE (essential MCU regulator), which was shown to be essential for Ca^2+^ uptake in metazoa, was affinity-purified from human cells in complex with MCU (*18*).

Even in the absence of structural data on the MCU complex, mitochondrial Ca^2+^ uptake and its regulation in mammals has been assumed to rely on a complex comprising MCU, EMRE, MICU1, and MICU2 (*10, 19-21*). Previous models generally believe that MICU1 and MICU2 form a cap to occlude the MCU channel in low [Ca^2+^] conditions, and when [Ca^2+^] is elevated, through conformational changes of the EF hands in these two regulators, they will depart from the MCU/EMRE pore to allow Ca^2+^ permeation (*22, 23*).

Several MCU structures from fungi and an MCU-EMRE supercomplex (MES) structure from human have been solved lately (*24-28*), however, no intact structure of the MCU-EMRE-MICU1-MICU2 supercomplex (MEMMS) has been reported. In the present study, after extensive optimization of expression and purification steps, we obtained the high-resolution cryo-EM structures of the human MES and MEMMS. The pore profiles of both structures are almost identical. Ca^2+^ is bound in the selectivity filter of both MES and MEMMS, and there is no pore obstruction in either of the structures. Therefore, we propose that both structures are in a conductive conformation. The MEMMS structure clearly demonstrates how MICU1 and MICU2 bind to MES. The two regulators apparently do not occlude the MCU channel. Instead, they form a bridge linking the two MCU pores through direct interactions with EMRE subunits. This finding is in striking contract to the generally accepted model in which MICUs cap and occlude the MCU channel on its cytosolic side. The accompanying paper also demonstrates that MICU subunits do not occlude the MCU pore. Rather, MICUs potentiate MCU activity as cytosolic [Ca^2+^] is elevated (*29*).

## Results

### Structure determination

The human MCU supercomplex was expressed in HEK 293F cells containing BacMam viruses for each of the genes *mcu, mcub, micu1, micu2*, and *emre*. After extensive optimization of reactants, we obtained an abundant amount of high quality MES and MEMMS protein samples, pulled-down by the C-terminally Strep-tagged EMRE in purification buffer with or without (EGTA) Ca^2+^, respectively. These samples (MES and MEMMS) were used to pool the grids for cryo-EM analyses (fig. S1, A and B, and Methods). Images were recorded with a combination of a Titan Krios Cryo-EM and a K2 direct electron detector in super-resolution mode (fig. S1, C and D). After extensive 2D and 3D classification of particles, a subset of particles was subjected to refinement, resulting in a 3D density map of MES at an overall resolution of 3.41 Å, and a density map of MEMMS at an overall resolution of 3.64 Å (Gold-standard FSC 0.143 criterion) (*30, 31*) (fig. S1, E and F, and fig. S2). Further subregion refinement with two different masks for the helical region and the N-terminal region of MES improved the resolution for these two regions to around 3.27 Å. Subregion refinement with masks for MICU1-MICU2, the helical region, and the N-terminal region of MEMMS further improved the resolution for these regions to 3.71, 3.30, and 3.39 Å, respectively (Gold-standard FSC 0.143 criterion) (*30, 31*) (fig. S2).

We subsequently obtained our high-resolution human MES and MEMMS structures based on a combination of structure docking and *de novo* modeling (*32, 33*). The well-resolved density maps allowed us to build structural models for almost all residues with their side chains (Fig. 1, A and B, and figs. S3 and S4). However, three sets of densities were not optimal for model building. The first is the density for the highly conserved C-terminal poly-D tail (EMRE^101-107^: EDDDDDD) of EMRE (*18*), the second is the conserved N-terminal poly-K (MICU1^99-102^: KKKK) of MICU1, and the third is the density for the conserved C-terminal helix (around 450-470) of MICU1 (fig. S5, A and B). The ambiguity of these densities might be owing to the double strep tag we added, and the flexibility of these regions.

**Fig. 1.**
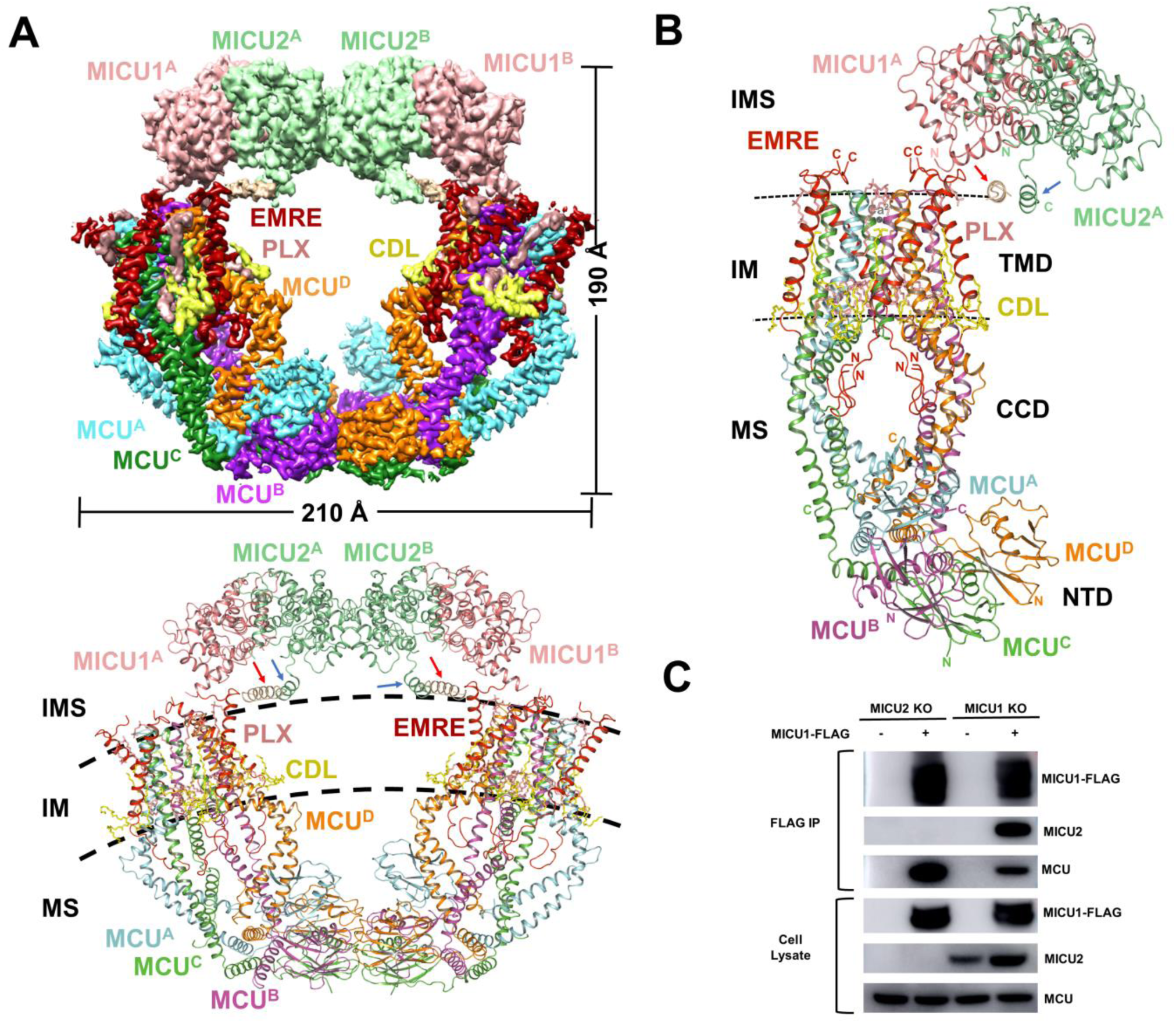
Overall structure of the MCU-EMRE-MICU1-MICU2 supercomplex. (**A**) Density map and structural model of MEMMS. PLXs, phosphatidylcholines, are shown in salmon; CDLs, cardiolipins, are shown in yellow; subunits of MEMMS are differently colored. IMS, intermembrane space; IM, inner membrane; MS, matrix. (**B**) Hetero-decamer of MEMMS. N and C termini of each subunit are labeled. The colors are the same as in A. The position of NTD, CCD and TMD of MCU are indicated. (**C**) FLAG co-immunoprecipitation of MICU1-FLAG expressed in MICU1 KO, MICU2 KO HEK 293T cells with transient expression of MICU1-FLAG. Lysates and elutes were immunoblotted with anti-FLAG, MCU or MICU2.

Despite our inclusion of a total of 5 transgenes in HEK 293F cells, the final structure only contains two kinds of subunits (MCU and EMRE) when the purification buffer contains Ca^2+^, and four kinds of subunits (MCU, EMRE, MICU1, and MICU2) when the expression and purification procedure is deprived of Ca^2+^ by adding EGTA. Within the MEMMS structure, we detected 2 Ca^2+^ ions, 8 cardiolipins (CDLs), and 16 phosphatidylcholines (PLXs). There were additional four Ca^2+^ ions within the MES structure (Fig. 1, A and B, and fig. S3, E and F).

### Overall structure of human MEMMS

Unlike previous fungal MCU homo-tetrameric structures (*24-27*), human MES forms a V-shaped structure comprising two hetero-octamers in C2 symmetry, in agreement with the human MES structure reported by Jiang’s team (*28*) (fig. S4A). Interestingly, human MEMMS forms an O-shaped ring, with a pair of MICU1-MICU2 heterodimer appearing like a bridge across the MES at the intermembrane space (IMS) side (Fig. 1A, and fig. S4B). When MICU1-FLAG plasmid was transfected into MICU2 KO HEK 293T cells, MICU1-FLAG was still able to co-precipitate with MCU, indicating that MICU2 is not required for interactions between MICU1 and MCU, which is consistent with our structure (Fig. 1C).

Alignment of MES and MEMMS shows that the helical region of these two complexes can perfectly match each other (fig. S4, D and E). Thus, in the later paragraphs, we mainly describe the structural features of MEMMS. MEMMS has a molecular weight of about 480 kDa and an overall dimension of 210 Å × 190 Å (Fig. 1A). The overall structure adopts the shape similar to that of two “goldfish,” as if glued together at both their heads (MICU1/MICU2 dimer) and tails (NTD of MCU), with the two transmembrane domains (TMDs) of MCU forming an angle reminiscent of ATP synthase dimers, located at cristae ridges of the inner mitochondrial membrane (*34-36*) (Fig. 1A).

The MCU subunit comprises three structural domains: the TMD, the coiled-coil domain (CCD), and the NTD (Fig. 1B). The TMD of MCU is known to be responsible for Ca^2+^ selectivity and conduction, and each MCU subunit contributes two transmembrane helices to the TMD: TM1 and TM2. Together, the four TM2s, which contain the highly conserved signature sequence (WDIMEP) (*13, 14, 17*), form the inner wall of the Ca^2+^ channel, and the four TM1s form the exterior wall of the channel. On the IMS side, TM1 and TM2 are linked by a short loop, forming a hairpin structure (fig. S6A). TM1 and TM2 are not parallel; instead, an obvious gap is formed between the two helices on the matrix side, which is filled by one PLX and one CDL molecules (Fig. 1B and fig. S6A).

The CCD and NTD of MCU are located in the mitochondrial matrix. The CCD of each MCU subunit comprises three α-helices: an exceptionally long and obviously bent helix (cc1), a lateral helix (cc2), and a short helix (cc3). Helix cc1 is extended from TM1, forming the coiled-coil structure in CCD with helix cc3. Helix cc2 links TM2 and cc3 (fig. S6A). The CCDs from human and fungi (*24-27*) both appear as “swollen bellies”. However, the human CCD is considerably larger than fungal ones. The NTDs of the four MCU subunits align in a configuration resembling that of bent “goldfish tails”. The NTD is connected to the cc1 via the linking helix α1, and the four α1 helices of each MCU subunits stably interact with each other, forming a four-helix bundle that stabilizes the MCU tetramers (fig. S6B).

### EMRE encages MCU and directly interacts with MICU1

EMRE is a small protein, containing only 107 residues after translation, and its 47 N-terminal residues are cut off after its transport into mitochondria (*18*). EMRE has been known to be required for mitochondrial Ca^2+^ uptake in human cells (*18, 37*). Within the membrane, four EMREs, four CDLs, four horizontal PLXs, and four vertical PLXs form a cage. They bundle up the four MCU subunits, thereby stabilizing and likely supporting a functional conformation of the Ca^2+^ channel (Fig. 2, A and B).

**Fig. 2.**
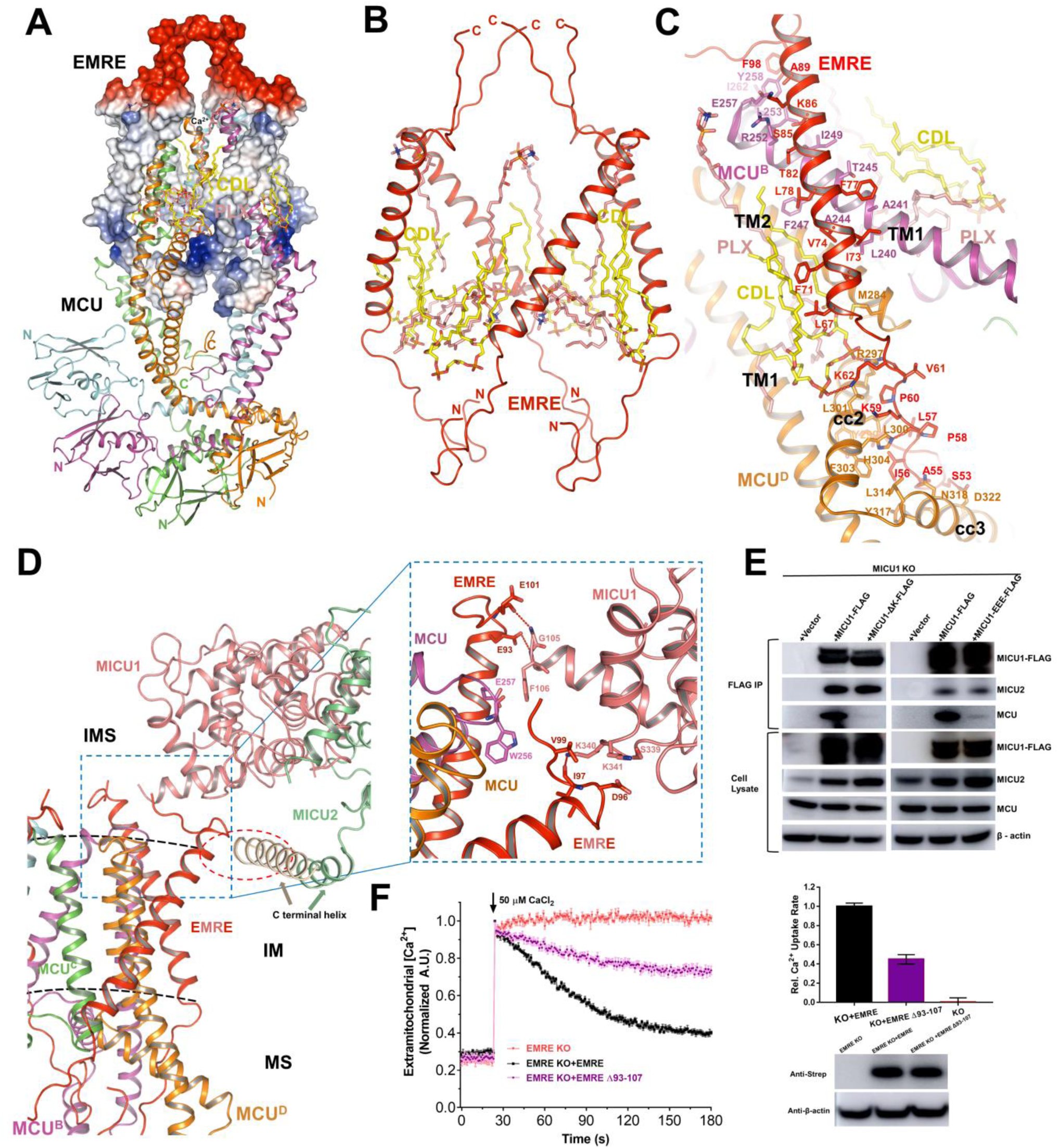
Interactions and functional roles of EMRE. **(A)** Four EMRE subunits form a cage surrounding the TMD of MCU. EMRE subunits are shown in an electrostatic surface model. The N-terminus of EMRE is positively charged, while the C-terminus is negatively charged. MCU subunits are shown in a cartoon model. Viewed from the membrane plane. **(B)** EMRE subunits and phospholipids form a cage that bundles up the central MCU tetramer. Four EMRE subunits are distinguished by red and all N and C-terminals are noted. PLXs are colored salmon. CDLs are colored yellow. (**C**) Detailed interaction between an EMRE (colored red) subunit and two MCU subunits (colored magenta and orange, respectively). Each EMRE interacts with two MCU. Transmembrane helix of EMRE interacts with TM1 of one MCU subunit, the N-terminal domain of EMRE interacts with the neighboring MCU cc2 and cc3. Residues responsible for interactions are labeled and shown as sticks. Hydrogen bonds are shown as red dashed lines. (**D**) Interactions between EMREs and one MICU1. The blue dashed box indicates the interactions between the tails of two EMRE and the one MICU1, the right enlarged dashed box shows the detail. Residues responsible for interactions are shown as sticks. Hydrogen bonds are shown as red dashed lines. The red dashed circle indicates interactions between MICU1 C-terminal helix and EMRE. (**E**) FLAG co-immunoprecipitation of MICU1-FLAG and related mutant constructs expressed in MICU1 knockout HEK 293T cells. Cells were transfected with MICU1-FLAG, MICU1 ΔK-FLAG or MICU1-S339E/K340E/K341E-FLAG plasmids (MICU1-EEE-FLAG). Lysates and elutes were immunoblotted with anti-FLAG, MCU, MICU2 or β-actin. Mean ± SEM, n ≥ 3 independent measurements. (**F**) The mitochondrial Ca^2+^ uptake of EMRE mutants at EMRE C-terminal. EMRE KO cells transiently expressing C-terminal strep tagged EMRE, or strep tagged EMRE Δ93-107 were given a ∼40 μM Ca^2+^ pulse. EMRE Δ93-107 (colored purple) suppresses channel function compared with wild-type EMRE (colored black) expressed in EMRE KO cells. Representative traces are shown on the left and bar graph in the right (mean ± SEM, n ≥ 3 independent measurements). Western blot of cell lysates from the different groups were performed to make sure that EMRE expression was similar by using antibody to strep. β-actin was used as the loading control.

In our structure, the N-terminal (residues 48-65) and C-terminal (residues 97-107) residues of EMRE appear as loops, while its middle residues (residues 66-96) adopt the configuration of a single α-helix. This middle EMRE α-helix locates within the membrane and is tilted by 37° relative to the membrane plane normal, such that each EMRE subunit interacts with two neighboring MCU subunits (Fig. 2, B and C). The N-terminal loop of EMRE protrudes into the particularly large chamber of CCD, forming rich hydrogen bonds with cc2 and cc3, and even with a CDL molecule (Fig. 2C).

The negatively charged C-terminal loops of EMRE protrude into the IMS and are responsible for direct interaction with MICU1. The structure shows no direct interaction between MICU1 and MCU. The positively charged N-terminal poly-K (residues 99-102) region of MICU1 and the negatively charged C-terminal tail (residues 93-107) of EMRE are in close proximity to each other, as shown by clear interactions between MICU1 Gly^105^, Phe^106^ and EMRE Glu^93^, Glu^101^ (Fig. 2D), consistent with previous functional study which has detected interactions between these two oppositely charged tails (*38*). To confirm the linking role of MICU1 poly-K region, we deleted these amino acids and subsequently found that MICU1-ΔK cannot co-precipitate with MCU (Fig. 2E). In addition, we found Ser^339^, Lys^340^, Lys^341^ sequence in MICU1 C-lobe that can also interact with the negatively charged tail of another adjacent EMRE (Fig. 2D). So, we introduced a triple mutation (S339E, K340E, K341E) in MICU1 and found that the triple mutant also has reduced interaction with MCU. Finally, we truncated the negatively charged tail (residues 93-107) of EMRE, and the Ca^2+^ uptake rate of MCU complex in high [Ca^2+^] condition was reduced about twofold (Fig. 2E). These results indicate that the N-terminal domain and Ser^339^, Lys^340^, Lys^341^ sequence of MICU1 are important for its recruitment onto MCU/EMRE complex, most probably through interactions with the negatively charged C-terminal tail of EMRE. Furthermore, the interaction between the MCU/EMRE pore and the MICU1/MICU2 dimer appears to have a positive effect on the Ca^2+^-transport activity.

### MICU1 and MICU2 do not occlude the MCU pore

MEMMS are linked at the IMS side via MICU1 and MICU2, and at the matrix side via MCU NTDs (Fig. 1A). Each MICU1 and MICU2 subunit contains four EF-hands, of which two are capable of binding Ca^2+^ ions (*39*). However, in our structure, no Ca^2+^ is bound to MICU1 or MICU2 subunits, which can be attributed to the deprivation of Ca^2+^ by EGTA during expression and purification (Fig. 3A). Alignment of MICU1 and MICU2 shows that they have very similar core structures (N-lobe and C-lobe), but their N-terminal domains and C-terminal helices are different (Fig. 3A). The N-terminal domain and Ser^339^, Lys^340^, Lys^341^ sequence of MICU1 can interact with EMRE as discussed above. MICU1 and MICU2 form a hetero-dimer in a previously reported ‘face-to-face’ pattern, while the two MICU2 subunits interact in a ‘back-to-back’ pattern (*40*). Consequently, the N and C lobes of MICU1 and MICU2 subunits are arranged in an alternative pattern to link two MCU channels (Fig. 3, B and C).

**Fig. 3.**
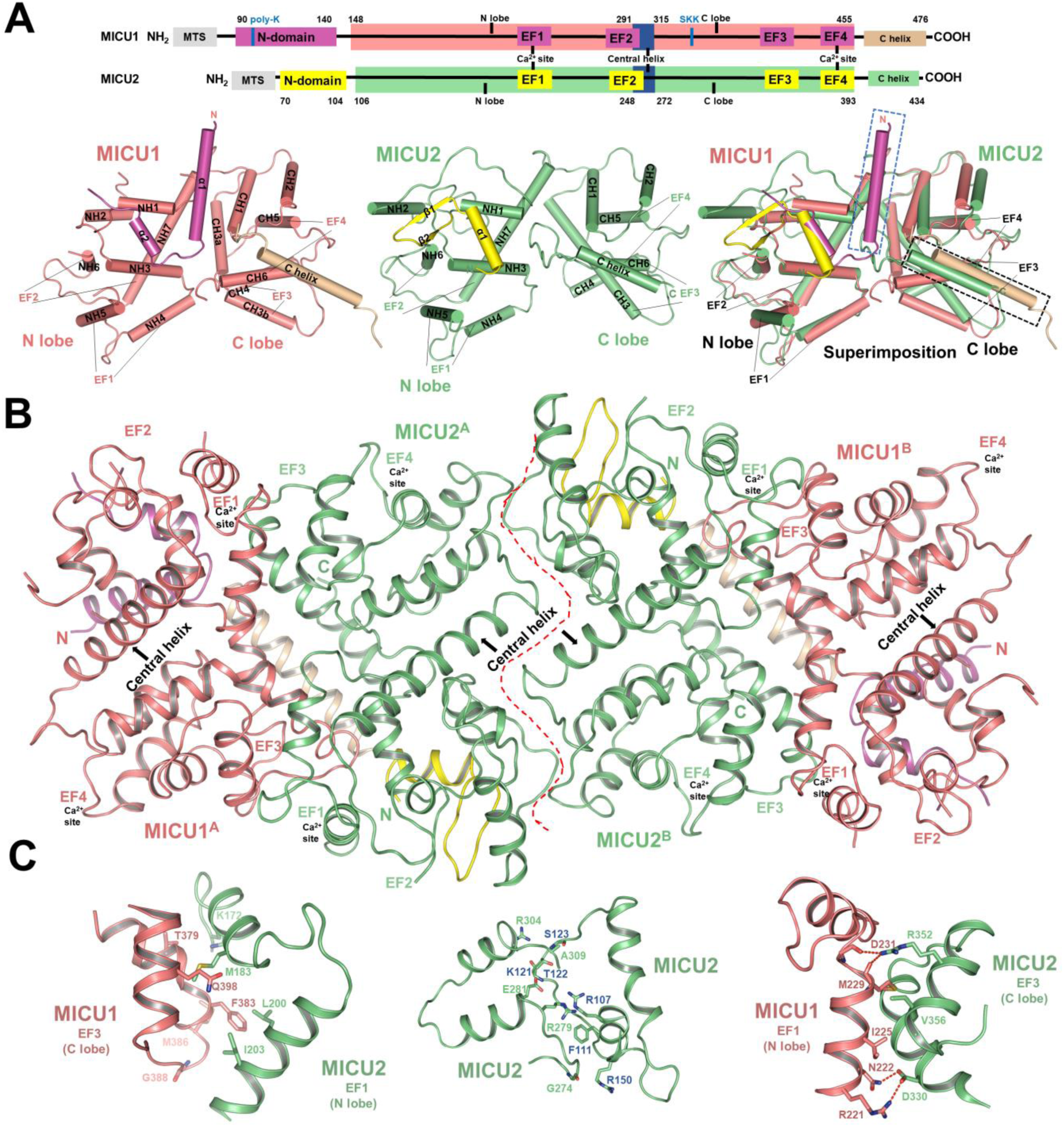
MICUs in the MEMMS. (**A**) Schematic domain organization (top), individual cartoon representation (lower left and middle panel) and superimposition (lower right) of the overall structure of MICU1 (the N-domain colored in magenta, the main body colored in light-pink and C-terminal helix colored in wheat) and MICU2 (the N-domain colored in yellow, the main body colored in light-green). The N-lobe, C-lobe, EF hand of each protein are indicated. In the superimposed MICU1 and MICU2 structure, the blue dashed box indicates the unique N-terminal helix of MICU1, the black dashed box indicates the C-terminal helix of MICU1. N or C termini of each MICU1 or MICU2 monomers are indicated. (**B**) The ‘face-to-face’ interaction between MICU1-MICU2 hetero-dimers and the ‘back-to-back’ interaction between two MICU2s. The colors are the same as in A, the central helix of each subunit is labeled. (**C**) Detailed interactions between MICU1 and MICU2. The left panel shows interactions between MICU1 C-terminal lobe and MICU2 N-terminal lobe. The middle panel shows interactions within MICU2 homodimer. The right panel shows interactions between MICU1 N-terminal lobe and MICU2 C-terminal lobe. Hydrogen bonds are indicated as red dash lines. The oxygen and nitrogen atoms are colored red and blue, respectively.

Previous models suggested that MICU1/2 dimer occludes the MCU pore (*22, 23, 39, 40*) at low cytosolic [Ca^2+^], which is obviously not the case as shown by the MEMMS structure (obtained in the presence of EGTA). The accompanying manuscript also reports that MICU subunits do not occlude the MCU channel in low cytosolic [Ca^2+^], using direct patch-clamp analysis of MCU currents (*29*). In addition to the EMRE/MICU1 interactions, the C-terminal helices of both MICU1 and MICU2 also contribute to MICU localization onto the inner membrane (fig. S7). In the MEMMS structure, although it’s difficult to analyze the detailed interactions between these two helices due to the vague local density, one can still appreciate that the two helices are parallel to each other at the surface of inner mitochondrial membrane (Fig. 2D and fig. S3G). The C-terminal helices of MICU2 have hydrophobic residues partially buried in the inner membrane, while the positively charged residues point parallel to the membrane, interacting with the negatively charged phosphates of the membrane (fig. S7). This is in agreement with the previous reports that MICU1 and MICU2 directly interact with the lipid membrane (*12, 41, 42*). Previous reports also show that the C-terminal helix is important for the interaction of MICU1 with MES (*43, 44*). Accordingly, deletion of MICU1 C-terminal helix significantly weakened the binding of MICU1 to MCU, and even lowered Ca^2+^ uptake activity (*43, 44*). Although the density map of this area is not clear enough for deciphering detailed interactions, we find that MICU1 C-terminal helix is in close vicinity of the EMRE helix (Fig. 2D). In a previous study, Co-IP assay showed that MICU2 ΔC could not interact with MICU1 or MCU (*39*). These findings are consistent with the MEMMS structure, in which the C-terminal helices act as an anchor to maintain MICU1 and MICU2 near to each other at the surface of inner mitochondrial membrane.

In the MES structure solved under high [Ca^2+^], no electron density can be associated with MICU1 or MICU2. We consider that the loss of MICUs is an artifact of the purification procedure. Since the membrane was solubilized in detergent, the C-terminal helix could lose its attachment with the membrane. In addition, when EF hands are occupied by Ca^2+^, conformational change of MICUs likely makes them more vulnerable to dissociation, leading to loss of MICU1 and MICU2 in the MES structure. Ca^2+^ uptake in high [Ca^2+^] condition was impaired in MICU1ΔC cells (*44*), which also suggests MICU1 is very likely attached to MCU in high [Ca^2+^]. In fact, in a previous report, interaction between MCU and MICU1 in high [Ca^2+^] was detected through co-IP (*44*).

In the matrix, the four NTDs align in a bent “fish-tail” configuration (fig. S8A). Note that this fish-tail alignment is in agreement with the previously reported MCU NTD crystal structure from human (*45, 46*), cryo-EM structure of the zebrafish MCU (*25*), and the recently published MES structure from human (*28*), but is quite distinct from the MCU structures from fungi (*24-27*) (fig. S8B). At the interface between the two fish-tails in MEMMS, three NTD pairs are tightly connected, while one set of NTDs are spared, which is the same case in MES (fig. S8C). Interestingly, the interaction patterns are not identical for all three pairs, and the only polar interaction that occurs commonly for all three pairs is between Asp^123^ and Arg^93^. We mutated Asp^123^ to Arg, or Arg^93^ to Asp, respectively, and found that both mutations have negligible influence on the Ca^2+^ uptake rate (fig. S8D). Thus, the other interactions between the three NTD pairs could still hold the two MCU channels together *in vivo* after these mutations.

### Phospholipids and the matrix gate

In both MES and MEMMS structures, one CDL and one horizontal PLX molecules inserts into the gap between TM1 and TM2 of each MCU subunit, and another vertical PLX molecule stands alongside each TM2. Notably, most of the lipid chains of the CDL and vertical PLX in our structure are parallel to the helices of TMD, while the lipid chains of the horizontal PLX are positioned horizontally in the membrane (Fig. 4A and fig. S6A). In a previous fungal structure (*24*), a horizontal PLX molecule was also found in the wall of the MCU channel. However, when we compare these two structures, the position of these two PLX molecules does not match.

**Fig. 4.**
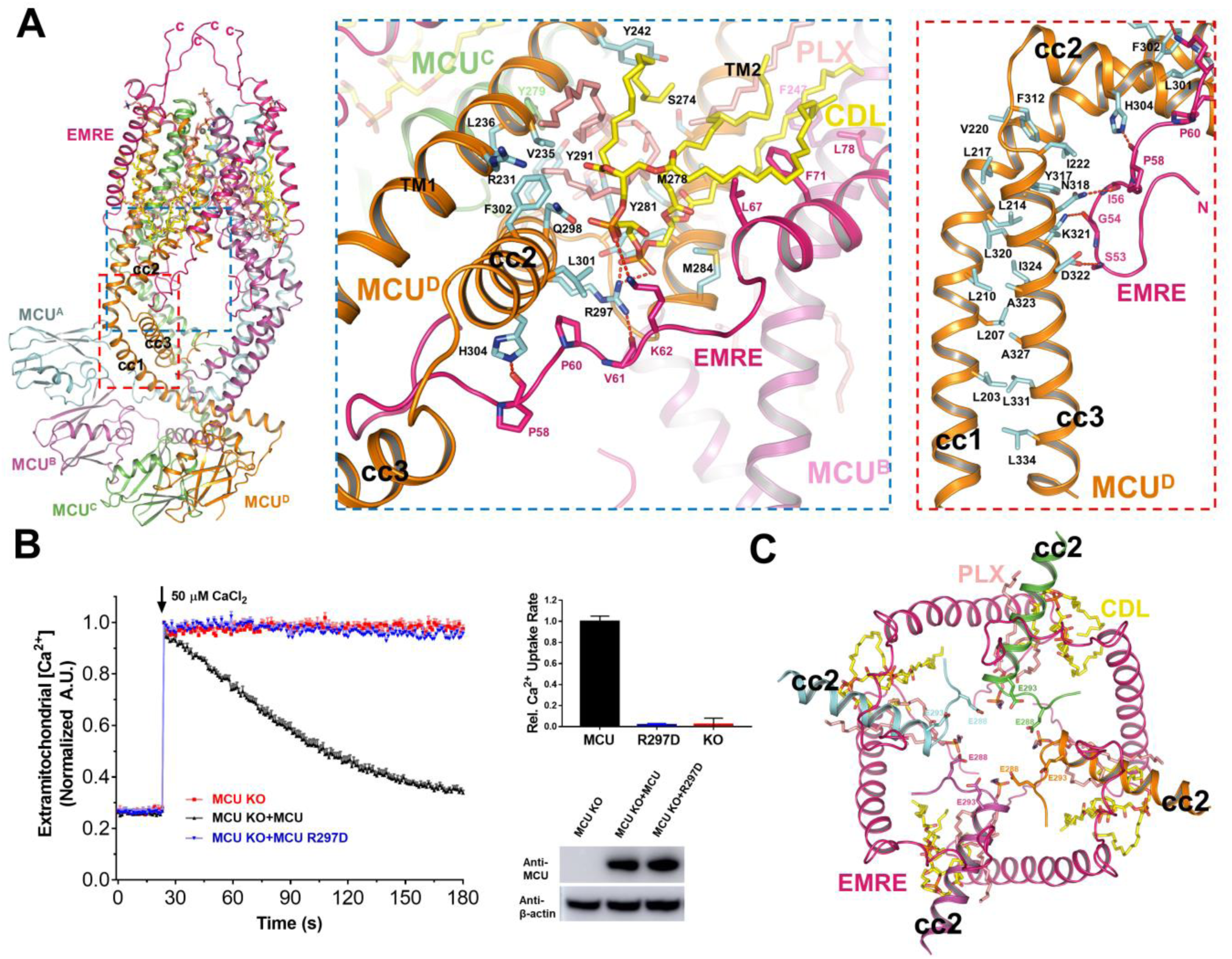
Interactions within the matrix gate of MCU. (**A**) Detailed interactions within the matrix gate of MCU complex. The blue dashed box indicates interactions between CDL and surrounding subunits, including two MCU subunits (colored magenta and orange, respectively) and one EMRE subunit (colored red), and interactions between the N-terminal of EMRE and cc2 of MCU. The red dashed box indicates the stable hydrophobic interface between MCU cc1 and cc3. Residues responsible for interactions are shown as sticks. Hydrogen bonds are shown as red dashed lines. (**B**) Mitochondrial Ca^2+^ uptake phenotype in R297D mutant of MCU. MCU KO cells transient expressing MCU, or MCU R297D were given a ∼40 μM Ca^2+^ pulse. R297D (colored blue) completely abolished the channel function. Representative traces are shown on the upper and bar graph in the bottom (mean ± SEM, n ≥ 3 independent measurements). Western blot of cell lysates from the different groups were performed to make sure the total amounts of protein expression were constant by using antibody to MCU. β-actin was used as the loading control. (**C**) Intrusion of the PLXs, CDLs and MCU cc2s into the central Ca^2+^ channel. Glu^288^s and Glu^293^s on cc2 are shown as sticks. EMREs, cc2, PLXs and CDLs are colored A.

In addition to interacting with TM1, TM2, and cc2 of one MCU subunit, each CDL molecule also interacts tightly with the neighboring EMRE. Specifically, a highly conserved residue of MCU cc2, Arg^297^, can form stable hydrogen bonds with both the phosphate group of CDL and the main chain oxygen of Val^61^ in EMRE (Fig. 4A), indicating Arg^297^ could be a critical residue for MCU regulation. This interaction (MCU-Arg^297^ to EMRE-Val^61^) is also observed in a previous human MES structure (*28*), however, interacting phospholipids were not detected. In the same report, authors truncated the N-terminal loop of EMRE, residue by residue until Lys^59^, and found that the Ca^2+^ uptake activity of MCU decreased gradually. Further truncation resulted in no EMRE expression, so the interaction between Arg^297^ and Val^61^ was not tested (*28*). To supplement, we mutated Arg^297^ to Asp, and strikingly this mutation completely abolished the Ca^2+^ uptake via MCU (Fig. 4B). Similarly, P60A mutation in EMRE, just next to the MCU-Arg^297^, can also totally abolish MCU activity (*47*), adding importance to correct interactions between cc2 and EMRE.

It has been proposed that cc2 and cc3 form a luminal gate near the matrix side of MCU that is maintained in an open conformation via its interaction with EMRE (*28, 48*). The previous human MES structure and our MES and MEMMS structures all detected stable hydrogen bonds between EMRE N-terminal loop and MCU cc2-cc3 (Fig. 4A), however, we also found several phospholipids filling the gaps between helices from MCU and EMRE. These phospholipids could stabilize the gaps and provide elasticity to this region, enabling the gate to be opened by EMRE (Fig. 4C). The MCU-R297D mutation might dissociate the bound CDL and disrupt the attachment of EMRE on cc2. This would leave cc2 free to roll aside and possibly push the negatively charged Glu^288^ and Glu^293^ residues of MCU inward, thus making the channel non-conducting. We observed multiple hydrophobic interactions between cc3 and cc1, which might help to achieve a correct position of the gate-forming cc2 (Fig. 4A and fig. S6A). The amino acid residues participating in these hydrophobic interactions are highly conserved and were shown to be indispensable for MCU activity (*48*). Besides, the negatively charged polar head of the horizontal PLX is also very likely involved in forming the gate, because their conformation is quite stable and they protrude deeply into the channel (Fig. 4C).

### The MCU pore and its regulation by MICU1/2

We detected three Ca^2+^ ions in each pore of the MES structure (using buffers containing Ca^2+^), and only a single Ca^2+^ ion in each pore in the MEMMS structure (using buffers deprived of Ca^2+^ by 0.1mM EGTA) (fig. S3, E and F). These Ca^2+^ ions are surrounded by the WDXXEP signature sequence (WDIMEP in human) adjacent to IMS, which was proposed to serve as a selectivity filter and discussed in detail previously (*24-28*). Four Asp^261^ residues form the first Ca^2+^ binding site, four Glu^264^ residues form the second, while four Tyr^268^ residues surround the third Ca^2+^ in MES (fig. S9, A and B). In the MEMMS structure, a single Ca^2+^ ion was detected at the high-affinity Glu^264^ site, the narrowest site in the pore (fig. S9C). The Glu^264^ site is likely the same site in the MCU pore that was previously reported to bind cytosolic Ca^2+^ with Kd ≤ 2 nM (*6*), and thus it should always be occupied by Ca^2+^ under physiological conditions. The lowest cytosolic [Ca^2+^] at which Ca^2+^ permeation through the MCU pore can occur is ∼100 nM (*22, 49, 50*). At this concentration, Ca^2+^ should start binding to the Asp^261^ site, which reduces affinity for Ca^2+^ binding at Glu^264^ and makes permeation possible. Our MEMMS structure does not have Ca^2+^ at the Asp^261^ site, and [Ca^2+^] in which MEMMS structure was determined was below 100 nM. This is expected, as the sample was washed multiple times with buffers containing 0.1 mM EGTA (see Methods). The estimated K_d_ for Ca^2+^ binding of MICU1/2 dimer is ∼600 nM (*51*), therefore it is also expected that the EF hands of MICU subunits in our MEMMS structure are Ca^2+^-free.

In MEMMS, MICU1/2 drag the two MCU tetramers closer to each other as compared to MES (fig. S4E). However, the pores of the MES and MEMMS structures are similar at both the selectivity filter and the putative matrix gate and show no possible obstructions for Ca^2+^ permeation (Fig. 5, A to C). This indicates that the binding of MICU1/2, does not occlude or obstruct the MCU pore. This conclusion is in a striking contrast to the currently accepted model of MICU1/MICU2 dimer (*22, 23, 39, 40*). However, our observations are consistent with the accompanying manuscript, in which authors also found that in low [Ca^2+^] conditions, the MCU pore is open and conducts similar currents in both the MICU1-deficient MCU complex (MES) and wide type MCU complex (MEMMS) (*29*).

**Fig. 5.**
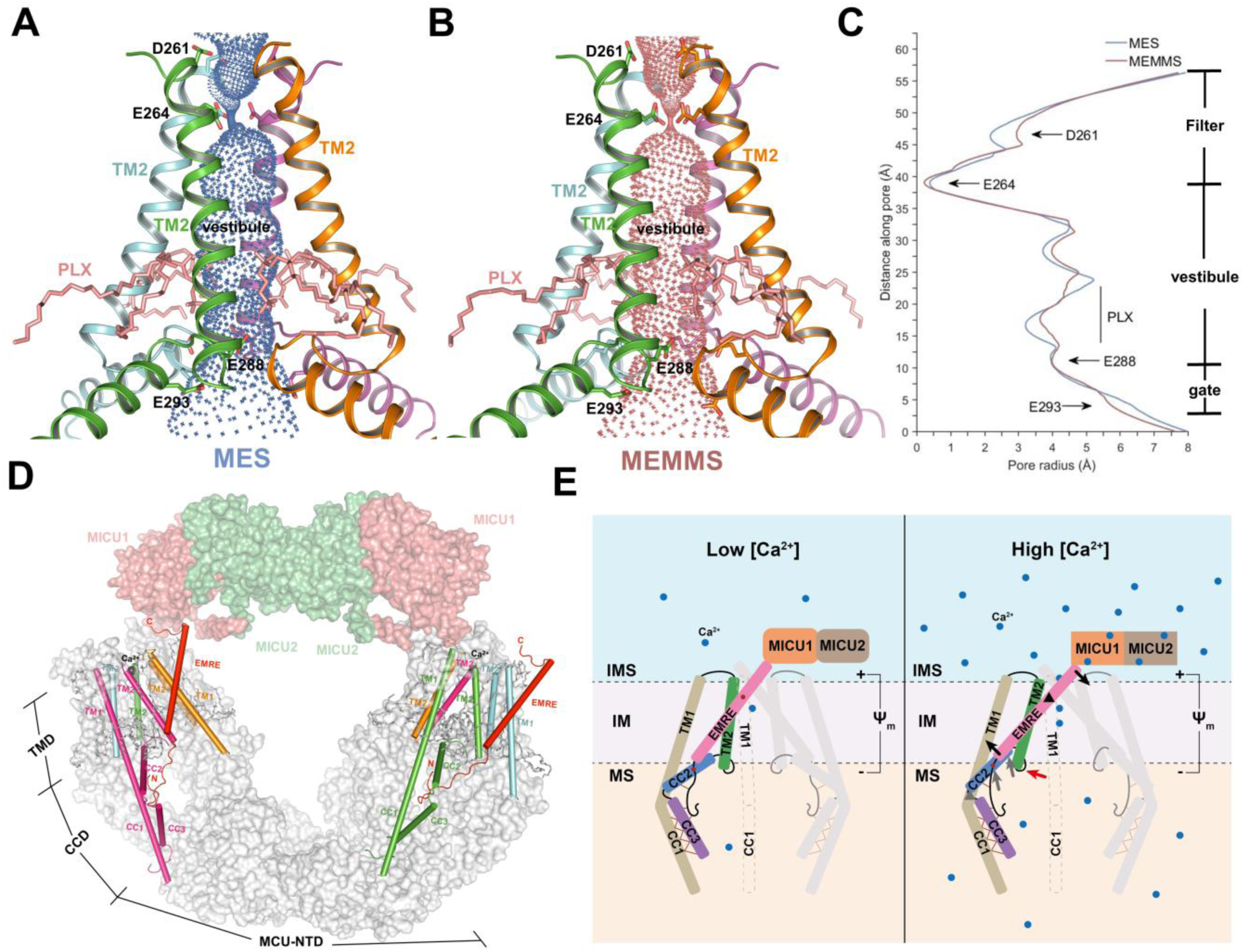
Comparison of pore profile in MES and MEMMS, and a proposed model of MCU complex regulation. (**A**) Cartoon model of the transmembrane pore of MES with the ion conduction pathway rendered in blue mesh. Asp^261^ and Glu^264^ at the entrance, and Glu^288^ and Glu^293^ at the exit of Ca^2+^ channel are shown as sticks. PLXs are shown as salmon sticks. (**B**) Cartoon model of the transmembrane pore of MEMMS with the ion conduction pathway rendered in brown mesh. Asp^261^ and Glu^264^ at the entrance, and Glu^288^ and Glu^293^ at the exit of Ca^2+^ channel are shown as sticks. PLXs are shown as salmon sticks. (**C**) Pore radius along the ion conduction pathway of MES and MEMMS. Filter, vestibule and gate are indicated, the gate residues and PLX are labeled. (**D**) EMRE anchors on the TM1 of an MCU, while the N-terminal interacts with the cc2 and cc3 of the neighboring MCU in the matrix, and the C-terminal interacts with MICU1 in the IMS, thus linking up MICU and MCU. All TM2 of MCU, typical TM1 and neighboring CCD domain are shown as cylindrical helices, the rest of MEMMS are shown as surface. (**E**) Proposed model of how EMRE and MICU regulate the conductivity of MCU supercomplex. Two sets of imagined levers are shown. EMRE is the first lever, with its pivot on TM1, its C-terminal loop attached to MICU1, and its N-terminal loop attached to CCD. cc2 is the second lever, with its pivot on the loop linking cc2 and cc3, its N-terminal attached to TM2, and its Arg^297^ attached to EMRE. Arg^297^ functions as the point of contact between the first and the second levers. Pivot and movement of the first lever is indicated by black triangle and arrows, respectively. Pivot and movement of the second lever is indicated by gray triangle and arrows, respectively. The movement of TM2 is marked by a red arrow. MICU1/2 conformational change is represented by a shape change. Membrane and membrane potential are labeled. The left panel is the low [Ca^2+^]; the right panel is the high [Ca^2+^]. IMS, intermembrane space; IM, inner membrane; MS, matrix.

With the four EF-hands of MICU1/2 dimers, MICU1/2 can sense the local cytosolic [Ca^2+^] in the vicinity of the MCU pore and undergo conformational change upon Ca^2+^ binding. The accompanying manuscript by Garg et al., demonstrates that Ca^2+^ binding to MICU1/2 potentiates Ca^2+^ permeation through the MCU pore by increasing the probability of its open state. Our MEMMS structure and the previously proposed MCU gating mechanism (*28*) could explain this functional behavior of the MCU complex. Specifically, we hypothesize that after Ca^2+^ binding to their EF-hands, a conformational change in MICU1/2 dimers exerts a force upon EMRE and the elastic MCU matrix gate, thus increasing its probability of open state.

## Discussion

In conclusion, here we report the first structure of intact MCU supercomplex as a 20-subunit O-shaped dimer of hetero-decamers, with auxiliary MICU1 and MICU2 subunits attached. We discovered that a pair of MICU1-MICU2 hetero-dimers link the two MCU channels, which is obviously different from previous models that assume MICU1/2 oligomers to ride across the MCU pore and occlude it in low cytosolic [Ca^2+^]. We found that MICU1 does not directly contact MCU, but can attach onto the MCU complex through interaction with EMRE, indicating that a critical function of EMRE is to couple the Ca^2+^-sensing MICUs with the MCU channel. We propose that upon Ca^2+^ binding to their EF hands, MICU1/2 exert a pulling force upon EMRE to stabilize the open state of the MCU matrix gate. These results are in agreement with the accompanying paper showing that Ca^2+^-free MICUs have no effect on ion permeation via MCU, and MICUs potentiate MCU function as cytosolic Ca^2+^ binds to their EF hands (*29*).

As shown in the MEMMS structure, two MICU1 and two MICU2 subunits form a straight line between the two MCU channels. EMRE could function like a lever, with its C-terminal loop interacting with MICU1, its central helix anchored to TM1 of MCU as the pivot, and its N-terminal loop supporting MCU CCD. The interaction between MCU Arg^297^ and EMRE Val^61^ is the force bearing point of cc2 (R297D mutation makes the MCU channel nonfunctional). In addition, we detected rich phospholipids around the MCU matrix gate (formed by the loop between TM2 and cc2), which could provide elasticity to this region. After Ca^2+^ binding, conformational change of MICU1/2 could exert a pull on the EMRE N-terminal, and cause the enlargement (or stabilization of the open state) of the MCU matrix gate (Fig. 5E).

We compared our MES structure with previous fungal MCU structures to find that human MES has a swollen CCD enlarged by EMRE (fig. S10). This curvature is very likely facilitated by Pro^216^ of cc1, which is conserved in mammals but absent in fungi (fig. S10E). The cc2s in reported fungal MCU structures are not well resolved, indicating that their position is flexible possibly due to lack of EMRE. The curvature of cc1 and the tight cc1-cc3 interaction could probably elevate the position of cc2 and close the gate if no EMRE is bound. Consequently, we propose that because fungal MCU does not have an elevated cc2, it does not require EMRE to maintain an open position. In contrast, EMRE is indispensable for human MCU because its cc2 is supported by EMRE N-terminal loop.

## Acknowledgements

We thank the Cryo-EM Facility Center of Southern University of Science & Technology (Shenzhen) and the Tsinghua University Branch of China National Center for Protein Sciences (Beijing) for providing facility support. We would like to thank Qingxi Ma for helpful editing of the manuscript. Computation was completed on the Yang lab GPU workstation.

## Author contributions

M.Y. conceived, designed, and supervised the project, built the model, analyzed the data and wrote the manuscript. W.Z., R.G., and L.Y. did the protein expression, purification, and detergent screening. H.Z., L.Z., and P.W. performed EM sample preparation, data collection and structural determination. W.Z., J.Y., Y.S. and W.Z. constructed the knockout cell lines, did the calcium uptake and Co-IP assays. All authors discussed the data of the manuscript.

## Data and materials availability

The atomic coordinates of the MES and MEMMS have been deposited in the Worldwide Protein Data Bank with the accession codes 6K7X and 6K7Y, respectively. The corresponding maps have been deposited in the Electron Microscopy Data Bank with the accession codes EMD-9944 and EMD-9945, respectively.

## Supplementary Materials

### Materials and Methods

#### Co-expression of human mitochondrial Ca^2+^ uniporter supercomplex

The optimized coding DNAs for *H*.*sapiens mcu* (Uniprot: Q8NE86), *mcub* (Uniprot: Q9NWR8), *micu1* (Uniprot: Q9BPX6), *micu2* (Uniprot: Q8IYU8) and *emre* (Uniprot: Q9H4I9) were synthesis in GenScript and cloned into the pEG BacMam vector (*51*), with tandem twin Strep-tag or FLAG tag at the C terminus of these five proteins. The BacMam viruses were produced and amplified in Sf9 cells. After extensively biochemical studies, we obtained the high quality and quantity protein complexes with the combination of the EMRE C-terminus tagged and the other four subunits without tags viruses. For larger amount protein purification, four liters of the HEK 293F cells were cultured. When the cell density reached 2×10^6^ cells per mL, the cells was transfected with the P3 (the third passage) BacMam viruses of MCU, MICU1, MICU2, MCUb and EMRE, each at 8 mL per liter cell culture. Transfected cells were cultured for 48 hours before harvesting. To capture the supercomplex in low [Ca^2+^], 0.1 mM EGTA was added into the medium and the cells were cultured for at least 1 hour before the same virus transfected.

#### Purification of the human mitochondrial Ca^2+^ uniporter supercomplex

All procedures were carried out at 4°C. To purify the MES complex, four liters of transfected cells were harvested, washed with 1×PBS and resuspended in 10 mM Tris pH 7.4, 225 mM sorbitol, 60 mM sucrose, 2 mM CaCl_2_ and 0.1% BSA, 1 mM PMSF. The suspension was homogenized by a soft blender for 150 s and the homogenate was centrifuged at 3,000 *g* for 10 min. Supernatant was further centrifuged at 20,000 *g* for 45 min to obtain the crude mitochondria. The pellet was suspended and extracted in 25 mM Tris pH 7.8, 150 mM NaCl, 2 mM CaCl_2_ with 1% digitonin. After incubation for an hour, the extraction was centrifuged at 20,000 *g* for 20 min at 4°C and the supernatant was applied to Strep-Tactin Sepharose by gravity at 4°C. The resin was washed three times with W buffer, which contained 25 mM Tris pH 7.8, 150 mM NaCl, 2 mM CaCl_2_ with 0.1% digitonin. The target proteins were eluted with W buffer plus 5 mM desthiobiotin, concentrated to 100 μL by 100 kDa cut-off centrifugal filter (Millipore) and further purified by Superdex200 increase 5/150 GL also in W buffer. To obtain the MEMMS complex, the same protocol was used with some modification: 2 mM CaCl_2_ in all the buffer was replaced by 0.1 mM EGTA. The peak fractions were collected for EM sample preparation, the presence of the complex was verified by Blue Native-PAGE and confirmed by mass spectrometry.

#### Sample preparation and cryo-EM data acquisition

4 μL aliquots of freshly purified MES or MEMMS at a concentration of 5 mg/mL were placed on glow-discharged 400-mesh Quantifoil R1.2/1.3 grids (Quantifoil, Micro Tools GmbH, Germany). Grids were blotted for 5 s and flash-frozen in liquid ethane using an FEI Mark IV Vitrobot operated at 8°C and 100% humidity. The grids were transferred to a Titan Krios (FEI) electron microscope equipped with a Cs corrector, operating at a voltage of 300 kV. Images were recorded by a K2 Summit direct electron detector (Gatan, Inc.) equipped with a GIF Quantum energy filter (slit width 20 eV) in the super-resolution counting mode. Data acquisition was performed using AutoEMation II with a nominal magnification of 105,000 times, which yields a super-resolution pixel size of 0.5455 Å on image plane, and with defocus ranging from −1.5 μm to −2.0 μm. The dose rate on the detector was ∼8.0 counts per pixel per second with a frame exposure time of 0.175 second and a total exposure time of 5.6 seconds. Each micrograph stack contains 32 frames. The total dose rate was approximately 50 e^-^/Å^2^ for each micrograph.

#### Image processing

A simplified flowchart of the procedure for image processing of MES is presented in fig. S2A. A total of 4,997 cryo-EM movie stacks were automated collected. The motion correction was performed using MotionCor2 (*52*) with 2×2 binning, resulting in a pixel size of 1.091 Å, and meanwhile, dose weighting was performed, yielding motion-corrected integrated images for further processing. After whole image CTF estimation using CTFFIND3, 4,706 good micrographs were manually selected from the dataset (*53*). A total of 471,401 particles were auto-picked using RELION-3.0 (*30*). After several rounds of two-dimensional (2D) classification, 327,716 particles were selected for further three-dimensional (3D) analysis. A total of 19,739 particles from the first 500 micrographs were used to generate initial models for the first round of 3D classification using RELION-3.0. Multi-reference 3D classification was performed for the 327,716 particles and 89,734 particles were selected then subjected to 3D refinement. Each particle was recentered using the in-plane translations measured in 3D refinement and re-extracted from the motion-corrected integrated micrographs. Gctf (*54*) was used to refine the local defocus parameters. The well centered particles with more accurate defocus parameters were subjected to 3D refinement without symmetry, which resulted in an electron density map at 3.67 Å resolution. The 3.67 Å map was fitted into a copy of itself rotated by 180° using Chimera (*55*), confirming a C2 symmetry in this map. Another round of 3D refinement with C2 symmetry was performed and yield a map at 3.51 Å resolution.

The 89,734 particles were further classified in to three classes using masked local angular search 3D classification with a step size of 0.9°and a local search range of 5°. Particles from two classes, which were in relatively good quality but slightly varied in the separation angle of the two MCU-EMRE hetero-octamer, were further refined and yield two maps at 4.18 Å and 3.41 Å resolution, respectively. To improve the density of N-terminal domains (NTDs), 3D refinement with a focused mask was performed for the 46,879 particles in the class of higher resolution, resulting in a focused map with a resolution of 3.27 Å after post processing. Since the density of the hetero-octamer transmembrane domain (TMD) and coiled-coil domain (CCD) of all the three classes could perfectly fit in each other, the 89,734 particles before the 3D classification were expanded according to C2 symmetry using relion_particle_symmetry_expand, then subtracted by the density of one NTD and the other whole hetero-octamer using RELION-3.0. The 179,468 subtracted particles were subjected to 3D refinement with a soft mask without symmetry, resulting in a focused map with a resolution of 3.27 Å after post processing. The focused map of NTDs and two copies of the focused map of TMD+CCD were fit into the 3.41 Å map then combined using PHENIX (*32*) Combine Focused Maps, resulting in the final map. The reported resolutions are based on the gold-standard Fourier shell correlation 0.143 criterion (*56*). All density maps were sharpened by applying a negative B-factor that was estimated using automated procedures (*57*). Local resolution variations were estimated using Resmap (*58*).

A simplified flowchart of the procedure for image processing of MEMMS is presented in fig. S2B. A total of 9,899 cryo-EM movie stacks were motion corrected, 2×2 binned and dose weighted using MotionCor2. After whole image CTF estimation using CTFFIND3, 9,113 good micrographs were manually selected from the dataset. A total of 986,805 particles were autopicked using RELION-3.0. After several rounds of 2D classification, 699,244 particles were selected and subjected to 3D classification, using the MES 3.41 Å map low-pass filtered to 50 Å as the initial model. After 3D classification, 250,977 particles of the class with an additional “cap” comparing to MES were selected then subjected to 3D refinement. Each particle was recentered and re-extracted from the motion-corrected integrated micrographs. Gctf was used to refine the local defocus parameters. The re-extracted particles were subjected to 3D refinement without symmetry, which resulted in a map at 4.09 Å resolution.

The 250,977 particles were further classified into three classes using masked skip-alignment 3D classification. A total of 45,864 particles of the class with clear “cap” were subjected to 3D refinement with a soft mask then yield a map at 3.64 Å resolution. Since the density of the MICUs (i.e. the “cap”) is still fragmentary, these particles were subtracted by the density of MES in the 3.64 Å map of MEMMS using RELION-3.0 and subjected to 3D refinement. Then 33,930 particles were selected after a final round of 3D classification and subjected to 3D refinement with a soft mask and C2 symmetry, leading to a reconstruction of MICUs at 3.71 Å resolution with much better density. To improve the density of NTDs, the 45,864 particles were subtracted by the density of CCDs, TMDs and MICUs, then subjected to 3D refinement with a soft mask and C2 symmetry, resulting in a density map of NTDs at 3.39 Å resolution. In order to improve the density of TMD+CCD, the 45,864 particles were expanded according to C2 symmetry using relion_particle_symmetry_expand, subtracted by the density of the rest part beside TMD+CCD, then subjected to 3D refinement with a soft mask. Eventually, the resolution of TMD+CCD was improved to 3.30 Å. The focused map of NTDs, MICUs and two copies of the focused map of TMD+CCD were fit into the 3.64 Å map then combined using PHENIX Combine Focused Maps, resulting in the final map.

The reported resolutions are based on the gold-standard Fourier shell correlation 0.143 criterion. All density maps were sharpened by applying a negative B-factor that was estimated using automated procedures. Local resolution variations were estimated using Resmap.

#### Model building and refinement and validation

The atomic model of MES was manually built and adjusted in COOT (*31*). And then, the model with the ligands were subjected to global refinement and minimization in real space refinement using PHENIX with secondary structure and NCS restraints. The crystal structure of MICU1 (PDB 4NSC) and MICU2 (PDB 6AGH) were used as the initial models for MICUs in MEMMS. The model was refined in real space using PHENIX with secondary structure and NCS restraints. The final atomic models were evaluated using MolProbity (*59*). Pore radii were calculated using the HOLE program (*60*). All the figures were prepared in PyMol (*61*).

#### Gene knockout by CRISPR/Cas9

Gene knockout by CRISPR/Cas9 was performed using a previously described protocol (*62*). Two sets of guide RNA sequences were designed. Guide sequences used for gene knockout were as follows: MCU-KO1: CAGGAGCGATCTACCTGCGG; MCU-KO2: TGAACTGACAGCGTTCACGC; EMRE-KO1: GGCTAGTATTGGCACCCGTC; EMRE-KO2: TACTAGCCAGCGAGCCGCTC; MICU1-KO1: AAACCAGTATGGGTATGCGC; MICU1-KO2: CGAATTTCAGCGTAAACTGC; MICU2-KO1: CAGCCGCGTCAGTGTTGCGG; MICU2-KO2: TGGGGCGGAAAACTGCGACG.

After sequencing, HEK 293T cells was transfected transiently with pSpCas9(BB)-2A-GFP plasmid which contained the corresponding guide sequence. Single cell was isolated by flow cytometry 24 hours later and proliferated for 2 weeks. Gene knockout was confirmed by sequencing and western blot.

#### Co-immunoprecipitation and western blot

All co-immunoprecipitation experiments were performed at 4°C. In brief, related HEK 293T knockout cells at 80%-90% confluence were transfected with 15 μg corresponding plasmids using lipofectamine 2000 (Thermo Fisher Scientific) and grown in a 37°C CO_2_ incubator for 24 hours. Transfected cells were lysed in 1 mL lysis buffer (25 mM Tris pH 7.8, 150 mM NaCl, 1 mM EGTA, cOmplete protease inhibitors) with 1% digitonin. The cell lysate was incubated for 30 min on ice and centrifuged for 10 min at 4°C at 20000 *g*. A small portion of the sample was used for whole cell lysate analysis and the rest was collected and incubated with anti-FLAG magnetic agarose (Thermo Fisher Scientific) for two hours in 4°C. The beads were collected on a magnet, washed three times with 1 mL lysis buffer which contained 0.1% digitonin, and eluted with 150 μL SDS-gel loading buffer for western blot.

For western blot analysis, proteins were subjected on a 4-20% SDS-PAGE gel (GenScript) and transferred onto a PVDF membrane (Millipore). Membranes were detected with the indicated antibodies. The primary antibodies were used: MCU (Abcam), MICU1 (Sigma-Aldrich), MICU2 (Sigma-Aldrich), FLAG (Easybio), Strep (Easybio), Mouse-β-actin (Easybio).

#### Mitochondrial Ca^2+^ uptake assays

Mitochondrial Ca^2+^ uptake was performed on MCU KO (MCU^-/-^), or EMRE KO (EMRE^-/-^) HEK 293T following the published protocol (*42*). Briefly, the transfected cells were digested, washed with 10 mL PBS for three times and re-suspended in buffer (25 mM HEPES pH 7.4, 125 mM KCl, 2 mM KH_2_PO_4_, 1 mM MgCl_2_, 10 μM EGTA, 5 mM glutamate, 5 mM malate, 3 μM thapsigargin, 0.005% digitonin, 1 μM Oregon Green-Bapta6F) to a final concentration of 10×10^6^ cells per mL. 150 μL cell suspension was transferred into a 96-well plate (Coring). Fluorescence was recorded using a Perkin/Elmer plate reader with excitation 488 nm/emission 535 nm before and after Ca^2+^ injection. 50 μM CaCl_2_ was injected, resulting in about 40 μM free Ca^2+^. The relative Ca^2+^ uptake rate is reported as the linear fit of the fluorescence for 3 minutes. In order to normalize the fluorescence readout, the maximal fluorescence value was set to 1.0 and the other fluorescence values at each time point were divided by the maximal fluorescence. The relative rate of Ca^2+^ uptake was analyzed in GraphPad Prism 7 (GraphPad Software, Inc.). Western blot was performed to ensure protein expression was comparable among the MCU or EMRE mutants in the uptake assay, Mouse anti-β-actin was used as a loading control.

**Fig. S1.**
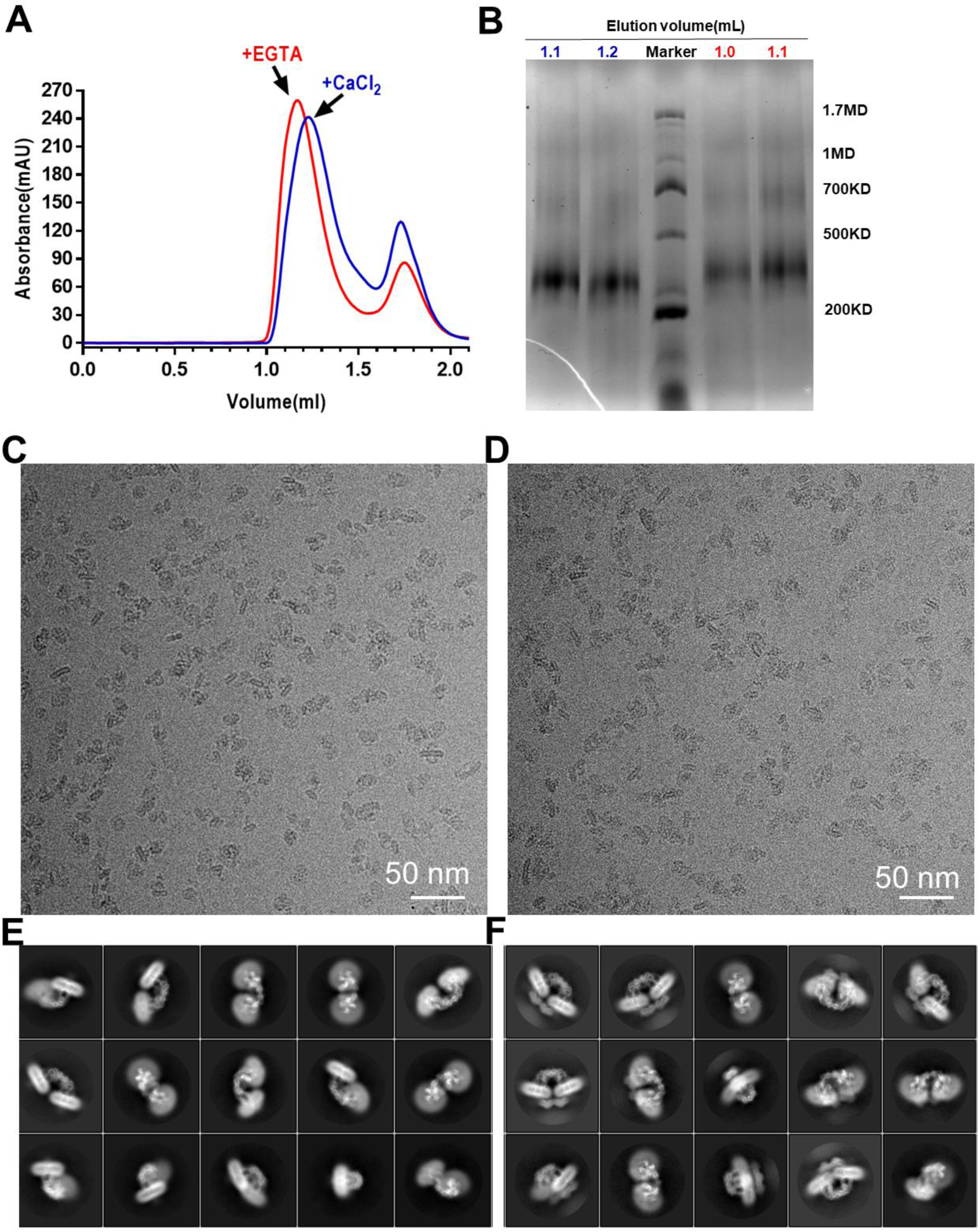
Structural characterization of MCU supercomplex. (A) Representative size-exclusion chromatography profile of the MEMMS (red trace) and MES (blue trace). (B) Protein samples of the MES and MEMMS size-exclusion chromatography fractions were subjected to BN-PAGE. Fractions of corresponded elution volume were used for cryo-EM sample preparation. (C) Representative micrograph of the MES. (D) Representative micrograph of the MEMMS. (E) 2-D class averages for the cryo-EM structure of MES. (F) 2-D class averages for the cryo-EM structure of MEMMS.

**Fig. S2.**
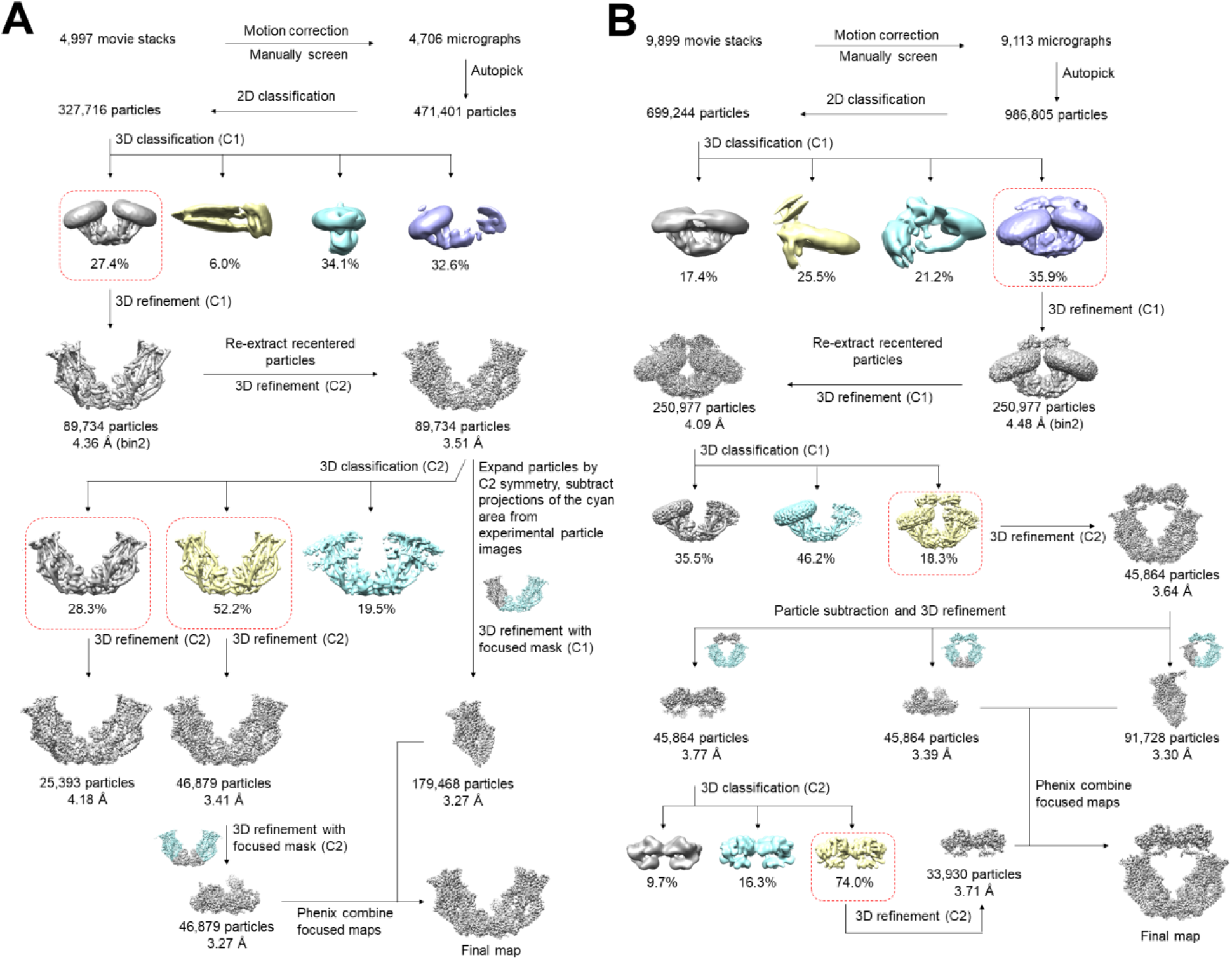
Flowchart of EM data processing of MES and MEMMS. Details of data processing are described in the ‘Image processing’ section of the Materials and Methods.

**Fig. S3.**
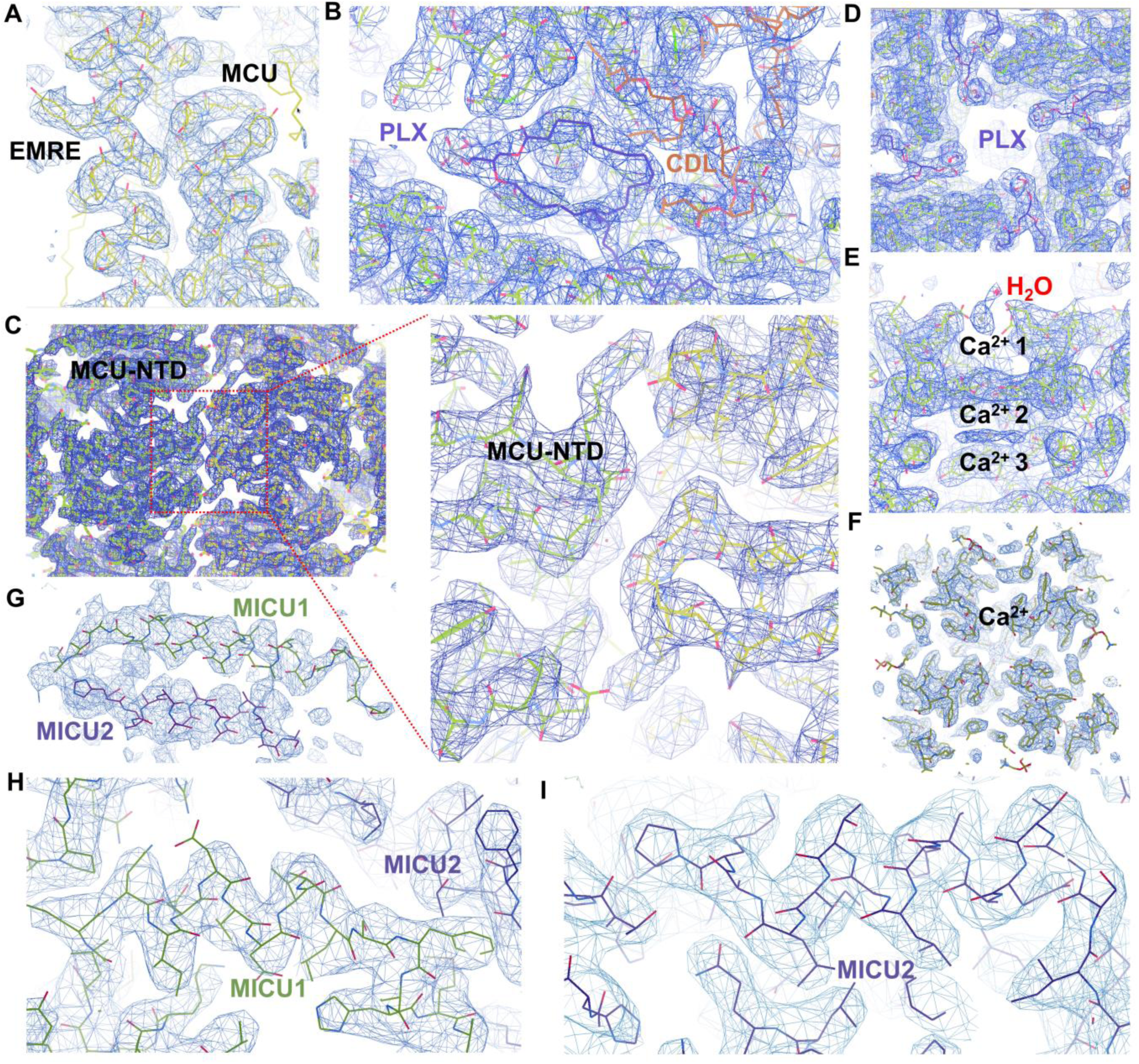
Representative density maps of MCU supercomplex. (A) Interaction between EMRE and MCU. (B) The electron densities of a PLX and a CDL. (C) Interaction between two MCU tetramers through three pairs of NTDs. (D) Polar heads of PLX intruding into central channel. (E) Entrance of central channel of MES. The density of three Ca^2+^ ions are shown. (F) Entrance of central channel of MEMMS. The density of one Ca^2+^ ion is shown. (G) C-terminal helix electron densities of MICU1 and MICU2. (H) Representative electron densities of interactions between MICU1 and MICU2. (I) Representative electron densities of MICU2.

**Fig. S4.**
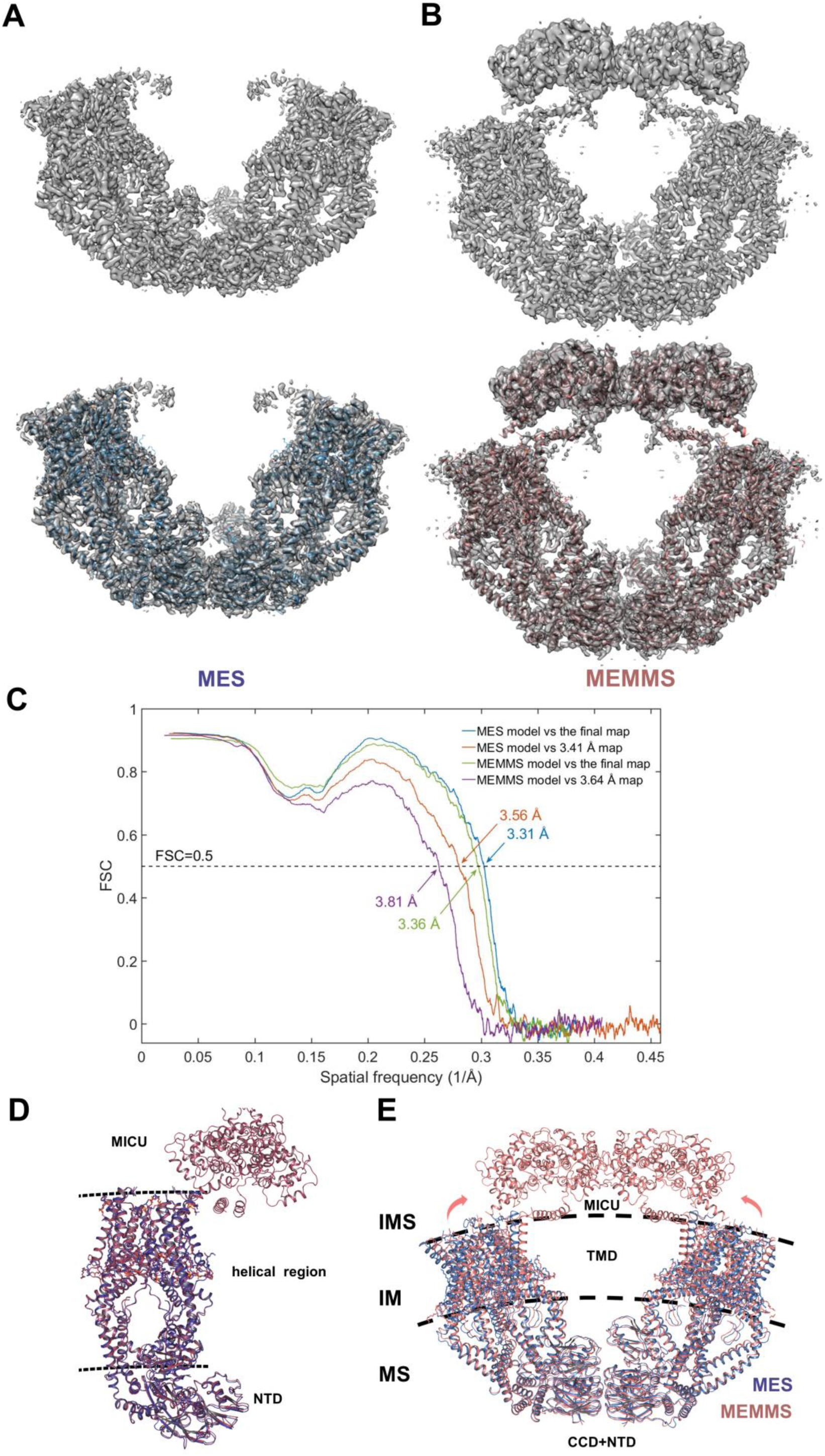
Model building of the MCU supercomplex. (A) Density map of MES viewed from membrane side (upper left). Merged density map and structural model of MES (lower left), model of MES is colored blue. (B) Density map of MEMMS viewed from membrane side (upper right). Merged density map and structural model of MEMMS (lower right), model of MEMMS is colored brown. (C) FSC curves of the refined model versus the final combined maps of MEMMS and MES that were refined against (blue) and the overall 3.64 Å and 3.41 Å map that were not refined against (orange), respectively. (D) Alignment of the MES octamer (blue) and the MEMMS decamer (brown). Membrane is indicated by two dashed lines. NTD, helical region and MICU is labeled. (E) Superimposition between MES and MEMMS. The overall structure of MEMMS is colored brown, and the overall structure of MES is colored blue. IMS, intermembrane space; IM, inner membrane; MS, matrix.

**Fig. S5.**
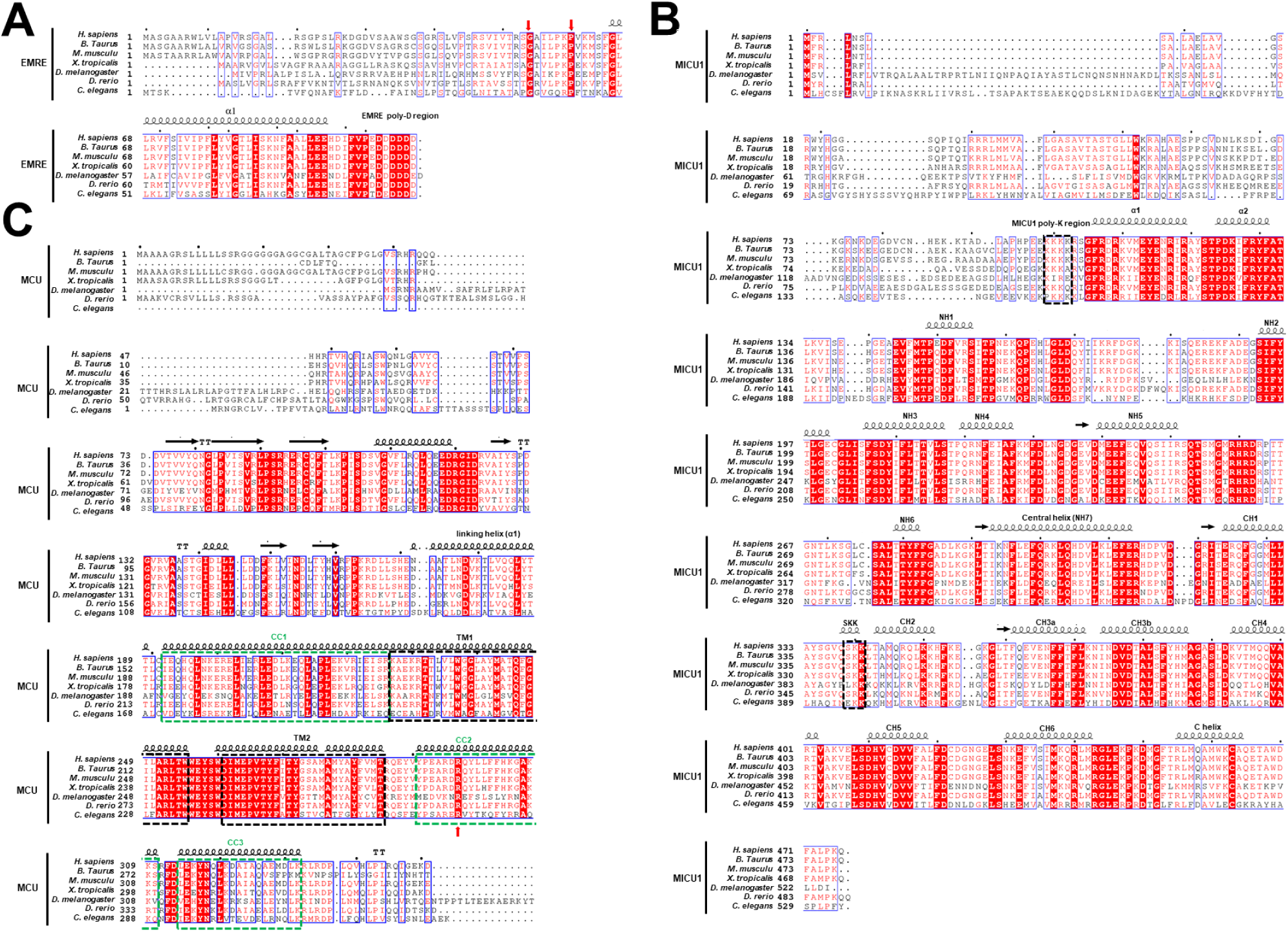
Structure-based EMRE, MICU1 and MCU orthologues alignment. (A) The amino acid sequences of *H. sapiens, B. taurus, M. musculus, X. tropicalis, D. melanogaster, D. rerio* and *C. elegans* EMRE are aligned and colored according to the ClustalW convention (Uniprot accession numbers: Q9H4I9, Q2M2S2, Q9DB10, Q28ED6, Q7JX57, A0A0J9YJ98 and Q9U3I4, respectively). Secondary structure represented by ribbons is based on the cryo-EM structure of *H. sapiens* EMRE. The conserved amino acid in the N terminal and the poly-aspartic tail are indicated. (B) The amino acid sequences of MICU1 are aligned and colored according to the ClustalW convention as in A. Secondary structure represented by ribbons is based on PDB 4NSC. The poly-lysine region and SKK region are indicated in dashed box. (C) The amino acid sequences of MCU are aligned and colored according to the ClustalW convention as in A. Secondary structure represented by ribbons is based on the cryo-EM structure of *H. sapiens* MCU. TM1 and TM2 are conserved and indicated in black dashed box. cc1, cc2 and cc3 domains are conserved and indicated in green dashed box. The conserved Arg in cc2 is indicated.

**Fig. S6.**
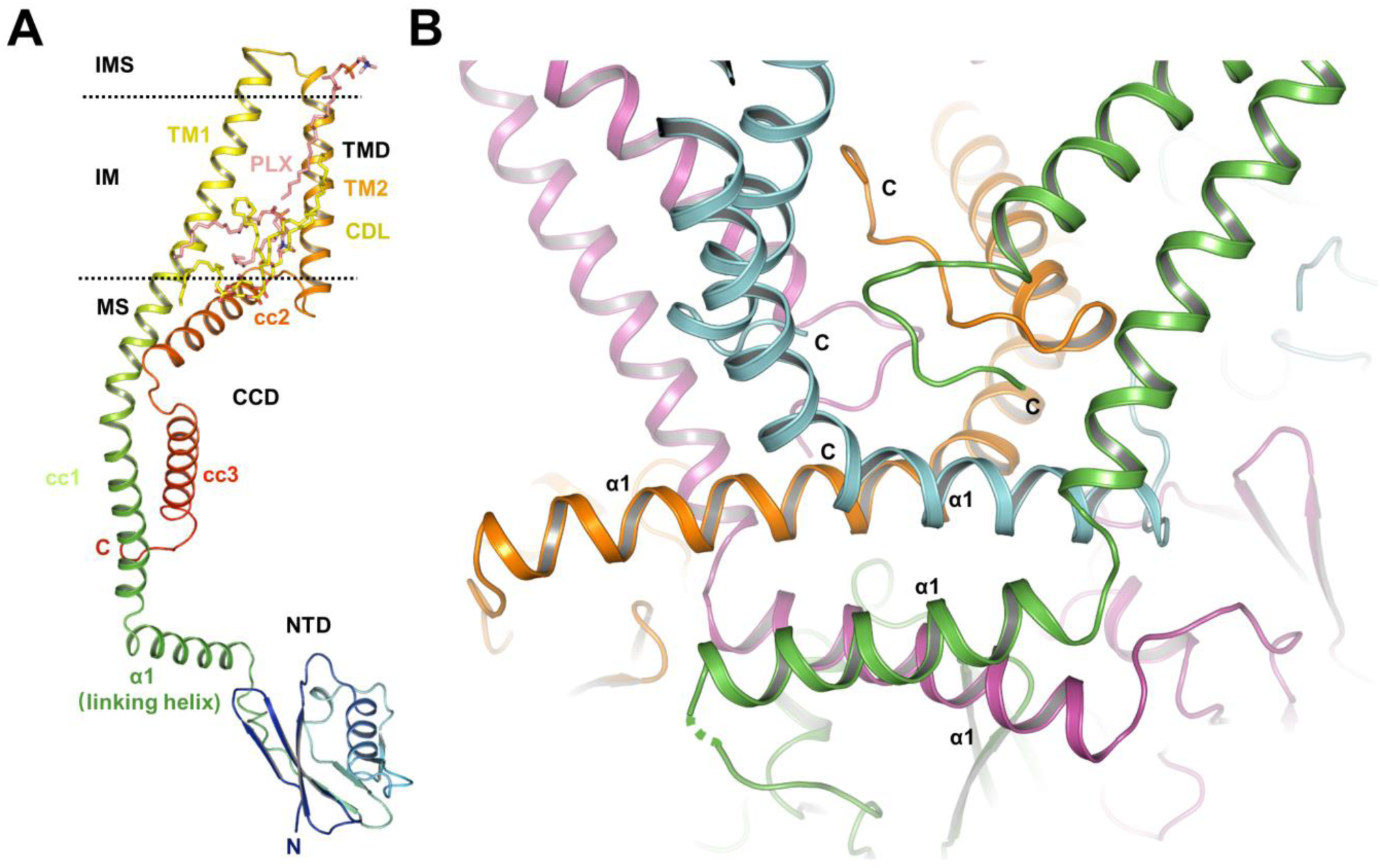
TMD, CCD and NTD of MCU. (A) Overall structure of the MCU monomer. Membrane is indicated by two dashed lines. TMD, transmembrane domain. CCD, coiled-coil domain. NTD, N-terminal domain. CDL and PLX are shown as sticks, colored in yellow and salmon respectively. MCU monomer is shown as cartoon and colored as rainbow. TM1 and TM2 form TMD. cc1, cc2, and cc3 form CCD. The linking helix α1 links CCD and NTD. (B) Linking site between CCDs and NTDs of MCU tetramer. Helices from four monomers are distinguished by different colors. C termini of different MCU monomers are indicated. Four α1s stack into two layers stabilizing the matrix region.

**Fig. S7.**
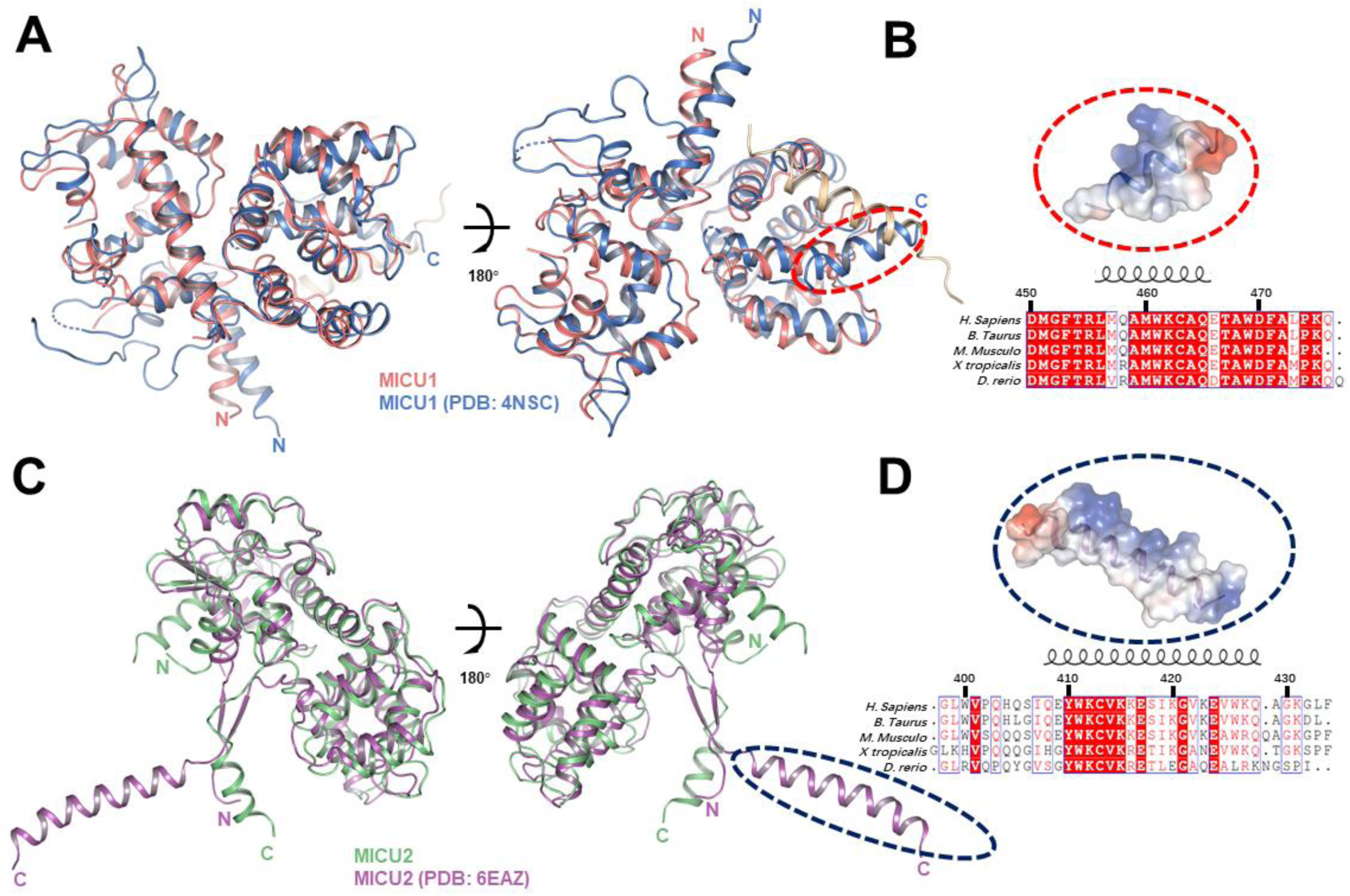
The C-terminal helix of MICU is important for MCU complex function. (A) Superimposition of MICU1 (colored in deep-salmon) in the MEMMS and hMICU1 in the Ca^2+^ free state (PDB 4NSC, colored in marine) in different views. 4NSC C-terminal helix is circled in red dashed line. (B) Surface electrostatic potential analysis of 4NSC C-terminal helix. The helix is positively charged on one side and hydrophobic on the other side. Amino acid sequences of *H*.*sapiens, B. taurus, M*.*musculus, X*.*tropicalis*, and *D*.*rerio* MICU1 C-terminal helix are aligned according to the ClustalW (Uniprot accession numbers: Q9H4I9, Q2M2S2, Q9DB10, Q28ED6, and A0A0J9YJ98, respectively). Secondary structure represented by ribbons is based on the C-terminal helix structure of 4NSC. Conserved residues among species are highlighted in red. (C) Superimposition of MICU2 (colored in lime-green) in the MEMMS and hMICU2 in the Ca^2+^ free state (PDB 6EAZ, colored in violet) in different views. 6EAZ C-terminal helix is circled in blue dashed line. (D) Surface electrostatic potential analysis of 6EAZ C-terminal helix. The helix is positively charged on one side and hydrophobic on the other side. Amino acid sequences of MICU2 C-terminal helix are aligned as in B. Secondary structure represented by ribbons is based on the C-terminal helix structure of 6EAZ. Conserved residues among species are highlighted in red.

**Fig. S8.**
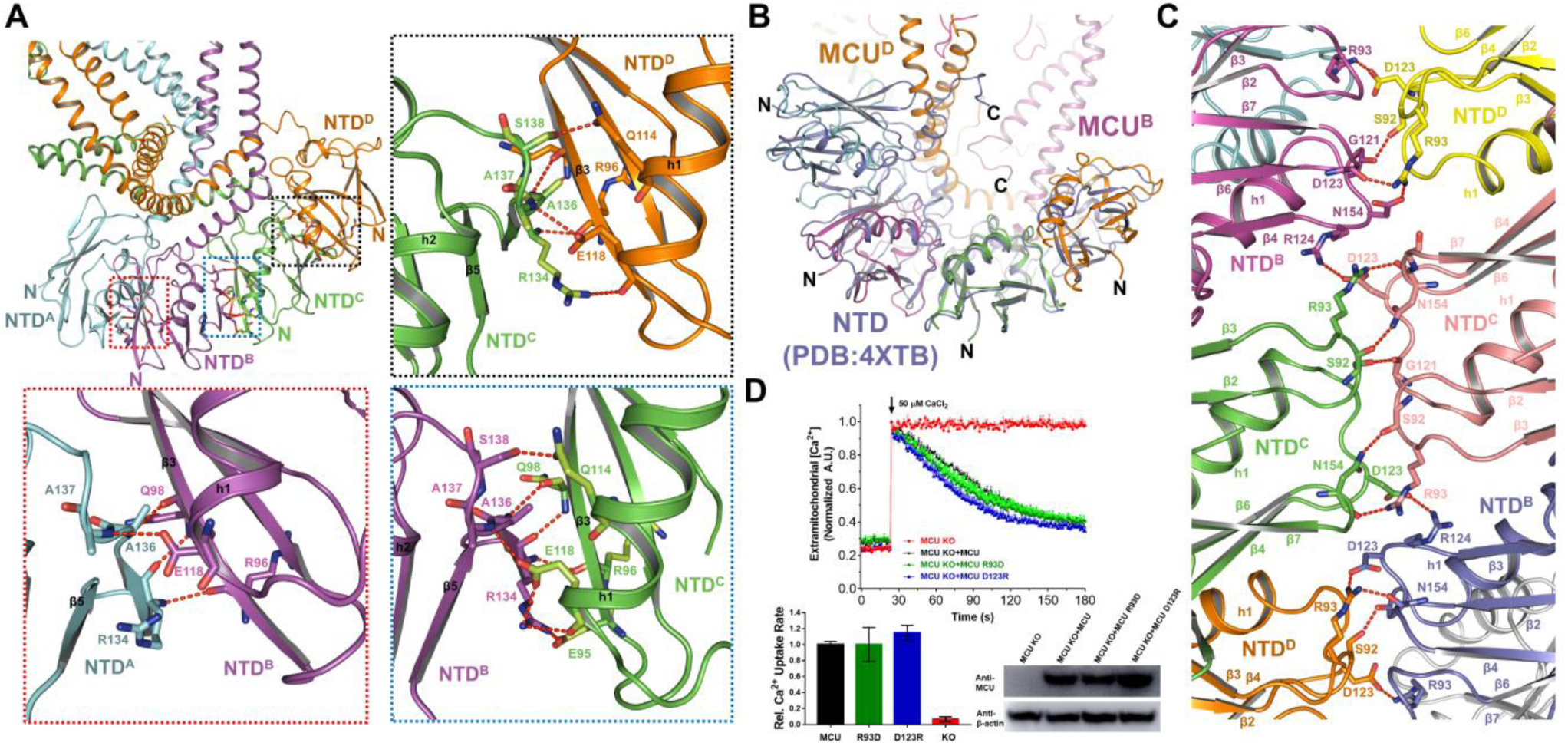
Interactions between NTDs of MCU in a crescent tail. (A) Four NTDs from the same MCU tetramer align to form a crescent tail. Four NTDs are distinguished by different colors, and labeled as NTD^A^, NTD^B^, NTD^C^, NTD^D^, respectively. N termini of MCU monomers are indicated. Three dashed boxes exaggerate three interacting sites between the four NTDs. Residues responsible for interaction are shown in stick. Hydrogen bonds are shown as red dashed lines. (B) Alignment of the four NTDs from our structure and the four symmetrical NTDs in the crystal lattice from the previous crystal structure of human MCU^NTD^ (PDB 4XTB). The NTD from the crystal structure is colored slate. The NTDs from different MCU subunits are distinguished by different colors. (C) Interaction between three pairs of NTDs from two MCU tetramers. Residues responsible for interactions are shown as sticks. Hydrogen bonds are shown as red dashed lines. (D) Mitochondrial Ca^2+^ uptake in permeabilized cells expressing MCU mutants based on MCU NTD interactions in MES/MEMMS structures. MCU KO cells expressing MCU, MCU R93D, or MCU D123R were given a ∼40 μM Ca^2+^ pulse. Representative traces are shown on the upper and bar graph in the bottom. Western blot of cell lysates from the different groups were performed to make sure the MCU expression was similar, using antibody to MCU. β-actin was used as the loading control.

**Fig. S9.**
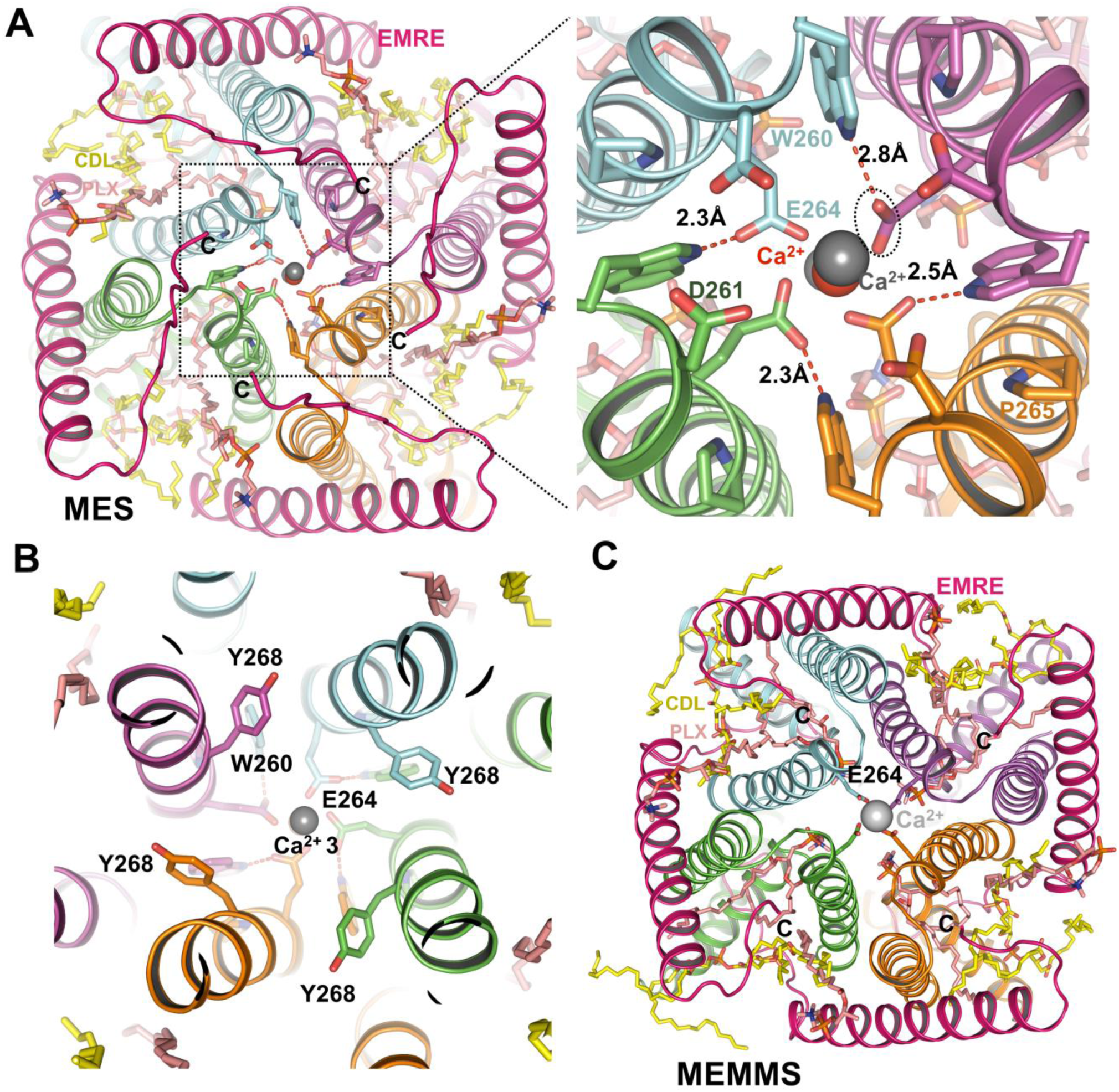
The entrance of Ca^2+^ channel of MES and MEMMS. (A) Ca^2+^ ion at the entrance of central channel of MES. The right panel is the zoomed-in image of left panel. The conserved residues in the selectivity filter are shown as sticks. Hydrogen bonds and their lengths are labelled. Ca^2+^ are shown as spheres. Ca^2+^ coordinating with Glu^264^ is shown as red sphere. (B) Helices from 4 different MCU monomers in MES are distinguished by different colors, encircling the central channel, as viewed from matrix side. Tyr^268^ is shown in stick point away from the central channel in our structure. Trp^260^ and Glu^264^ are also shown in stick. The third Ca^2+^ in the central channel are shown in sphere. (C)The sole Ca^2+^ ion coordinated by Glu^264^ residues is shown in the MEMMS structure, as viewed from the cytosolic side.

**Fig. S10.**
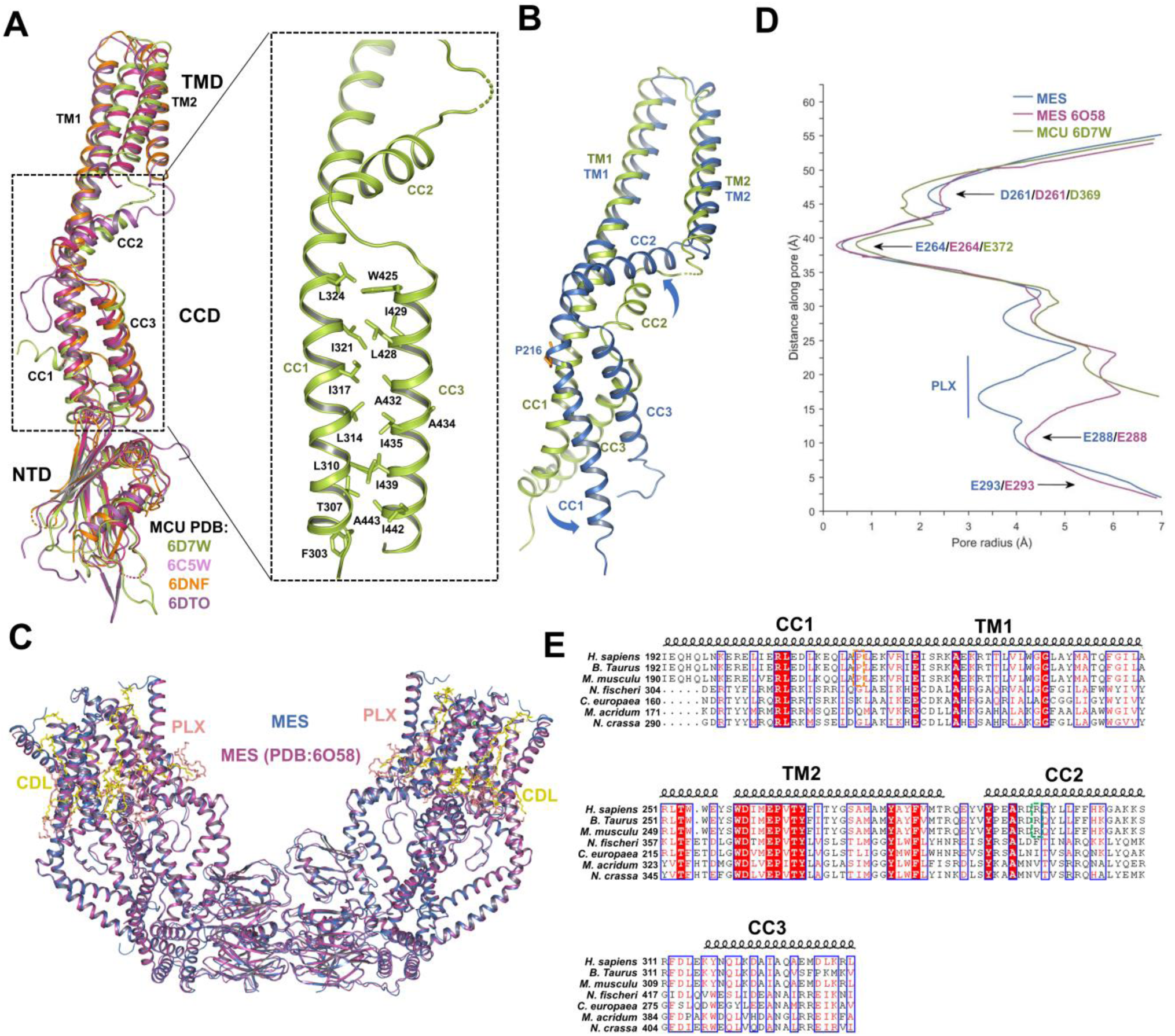
Comparison between two human MES and fungi MCU structures. (A) Superimposition of four fungal MCU structures (PDB 6D7W colored limon, 6C5W colored magenta, 6DNF colored orange, and 6DTO colored hot-pink). TM1 and TM2 in the transmembrane domain (TMD), cc1, cc2 and cc3 in coiled-coil domain (CCD), and NTD domain are indicated. The black dashed box shows that the interactions in CCD are conserved. The enlarged black dashed box shows the hydrophobic interactions within cc1 and cc3. (B) Alignment of *N*. *fischeri* MCU TMD+CCD domain (PDB 6D7W, colored limon) and *H*.*sapiens* MCU TMD+CCD domain (colored blue) based on TMD. TM1, TM2, cc1, cc2, and cc3 helixes are indicated. The blue arrows indicate the rotate directions of *H*.*sapiens* cc1, cc2, compared to *N*. *fischeri* cc1, cc2, respectively. Pro^216^ in *H*.*sapiens* cc1 is shown in sticks. (C) Superimposition of MES structure (colored blue) and MCU+EMRE structure (PDB 6O58) (colored purple). PLX and CDL are colored salmon and yellow, respectively. (D) Pore radius along the ion conduction pathway of the indicated MES or MCU structure. The gate residues and PLX are labeled. (E) The amino acid sequences of *H. sapiens, B. taurus, M. musculus, N. fischeri, C. europae, M. acridum* and *N. crassa* TMD+CCD are aligned and colored according to the ClustalW convention (Uniprot accession numbers: Q9H4I9, Q2M2S2, Q9DB10, A1CWT6, W2SDE2, E9DVV4 and Q7S4I4, respectively). Secondary structure represented by ribbons is based on the cryo-EM structure of *H. sapiens* TMD+CCD. The orange and green dashed boxes indicate the conserved Pro in cc1 and Arg in cc2 of higher eukaryotes, respectively.

**Table S1.**
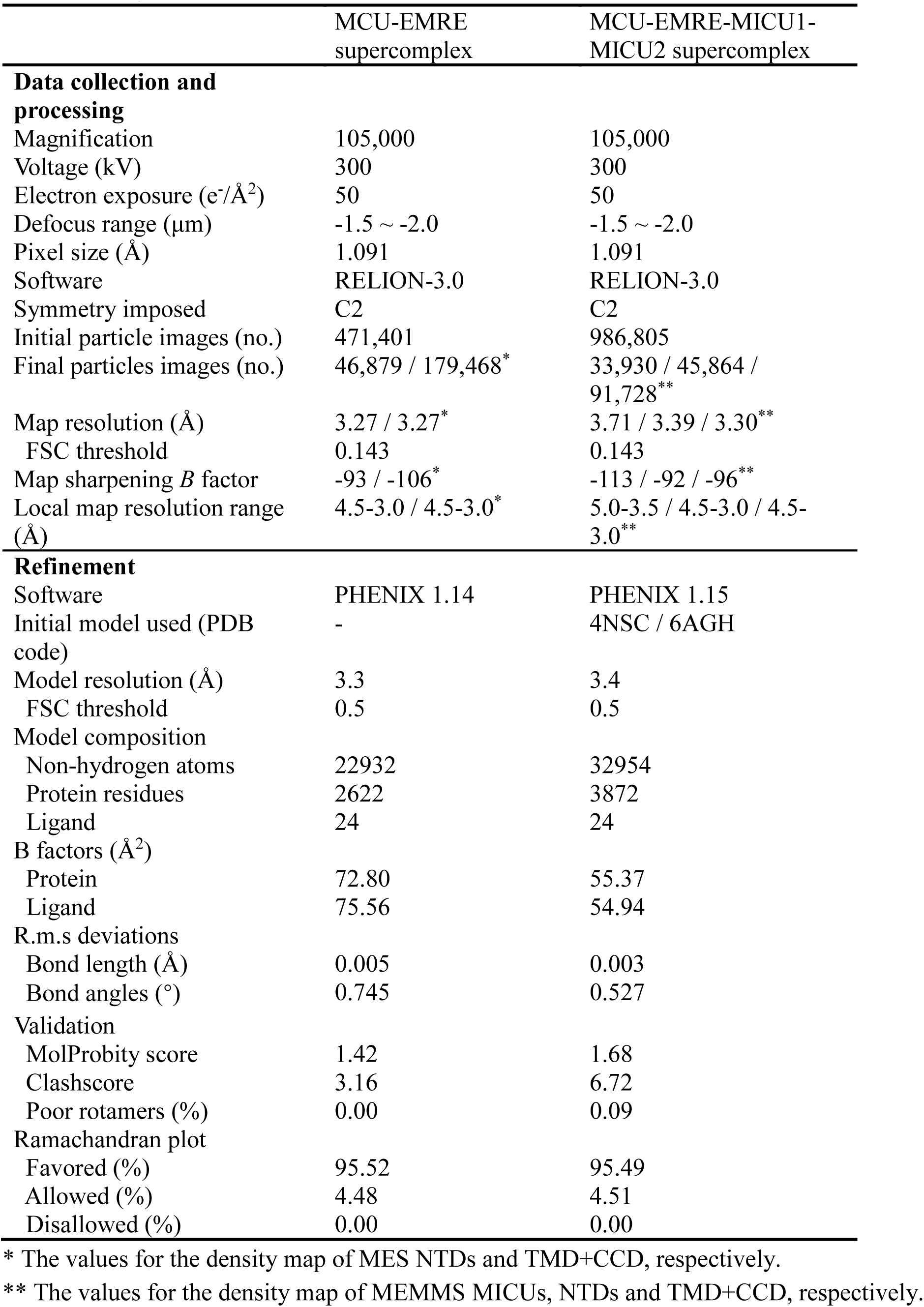
Cryo-EM data collection, refinement and validation statistics.

## References and notes

1. M. Gemba, E. Nakatani, M. Teramoto, S. Nakano, Effect of cisplatin on calcium uptake by rat kidney cortical mitochondria. Toxicol Lett 38, 291–297 (1987).

2. F. R. Mraz, Calcium and strontium uptake by rat liver and kidney mitochondria. Proc Soc Exp Biol Med 111, 429–431 (1962).

3. H. F. Deluca, G. W. Engstrom, Calcium uptake by rat kidney mitochondria. Proc Natl Acad Sci U S A 47, 1744–1750 (1961).

4. T. E. Gunter, D. R. Pfeiffer, Mechanisms by Which Mitochondria Transport Calcium. Am J Physiol 258, C755–C786 (1990).

5. F. D. Vasington, J. V. Murphy, Ca++ Uptake by Rat Kidney Mitochondria and Its Dependence on Respiration and Phosphorylation. J Biol Chem 237, 2670-& (1962).

6. Y. Kirichok, G. Krapivinsky, D. E. Clapham, The mitochondrial calcium uniporter is a highly selective ion channel. Nature 427, 360–364 (2004).

7. D. W. Jiang, L. L. Zhao, D. E. Clapham, Genome-Wide RNAi Screen Identifies Letm1 as a Mitochondrial Ca2+/H+ Antiporter. Science 326, 144–147 (2009).

8. R. Palty et al., NCLX is an essential component of mitochondrial Na+/Ca2+ exchange. P Natl Acad Sci USA 107, 436–441 (2010).

9. K. J. Kamer, V. K. Mootha, The molecular era of the mitochondrial calcium uniporter. Nat Rev Mol Cell Biol 16, 545–553 (2015).

10. D. De Stefani, R. Rizzuto, T. Pozzan, Enjoy the Trip: Calcium in Mitochondria Back and Forth. Annu Rev Biochem 85, 161–192 (2016).

11. M. Patron et al., The mitochondrial calcium uniporter (MCU): molecular identity and physiological roles. J Biol Chem 288, 10750–10758 (2013).

12. F. Perocchi et al., MICU1 encodes a mitochondrial EF hand protein required for Ca2+ uptake. Nature 467, 291–U267 (2010).

13. J. M. Baughman et al., Integrative genomics identifies MCU as an essential component of the mitochondrial calcium uniporter. Nature 476, 341–U111 (2011).

14. D. De Stefani, A. Raffaello, E. Teardo, I. Szabo, R. Rizzuto, A forty-kilodalton protein of the inner membrane is the mitochondrial calcium uniporter. Nature 476, 336–U104 (2011).

15. D. Chaudhuri, Y. Sancak, V. K. Mootha, D. E. Clapham, MCU encodes the pore conducting mitochondrial calcium currents. Elife 2, e00704 (2013).

16. M. Plovanich et al., MICU2, a Paralog of MICU1, Resides within the Mitochondrial Uniporter Complex to Regulate Calcium Handling. Plos One 8, (2013).

17. A. Raffaello et al., The mitochondrial calcium uniporter is a multimer that can include a dominantnegative pore-forming subunit. EMBO J 32, 2362–2376 (2013).

18. Y. Sancak et al., EMRE Is an Essential Component of the Mitochondrial Calcium Uniporter Complex. Science 342, 1379–1382 (2013).

19. N. Demaurex, M. Rosselin, Redox Control of Mitochondrial Calcium Uptake. Mol Cell 65, 961–962 (2017).

20. M. Ahuja, S. Muallem, The gatekeepers of mitochondrial calcium influx: MICU1 and MICU2. Embo Rep 15, 205–206 (2014).

21. A. G. Bick et al., Cardiovascular homeostasis dependence on MICU2, a regulatory subunit of the mitochondrial calcium uniporter. Proc Natl Acad Sci U S A 114, E9096–E9104 (2017).

22. M. Paillard et al., MICU1 Interacts with the D-Ring of the MCU Pore to Control Its Ca(2+) Flux and Sensitivity to Ru360. Mol Cell 72, 778–785 e773 (2018).

23. C. B. Phillips, C. W. Tsai, M. F. Tsai, The conserved aspartate ring of MCU mediates MICU1 binding and regulation in the mitochondrial calcium uniporter complex. Elife 8, (2019).

24. R. Baradaran, C. Wang, A. F. Siliciano, S. B. Long, Cryo-EM structures of fungal and metazoan mitochondrial calcium uniporters. Nature 559, 580–584 (2018).

25. C. Fan et al., X-ray and cryo-EM structures of the mitochondrial calcium uniporter. Nature 559, 575–579 (2018).

26. N. X. Nguyen et al., Cryo-EM structure of a fungal mitochondrial calcium uniporter. Nature 559, 570–574 (2018).

27. J. Yoo et al., Cryo-EM structure of a mitochondrial calcium uniporter. Science 361, 506-+ (2018).

28. Y. Wang et al., Structural Mechanism of EMRE-Dependent Gating of the Human Mitochondrial Calcium Uniporter. Cell 177, 1252–1261 e1213 (2019).

29. V. Garg et al., The Mechanism of MICU-Dependent Gating of the Mitochondrial Ca2+ Uniporter. bioRxiv, 2020.2004.2004.025833 (2020).

30. W. Du et al., Kinesin 1 Drives Autolysosome Tubulation. Dev Cell 37, 326–336 (2016).

31. J. Zivanov et al., New tools for automated high-resolution cryo-EM structure determination in RELION-3. Elife 7, (2018).

32. P. Emsley, B. Lohkamp, W. G. Scott, K. Cowtan, Features and development of Coot. Acta Crystallogr D Biol Crystallogr 66, 486–501 (2010).

33. P. D. Adams et al., PHENIX: a comprehensive Python-based system for macromolecular structure solution. Acta Crystallogr D Biol Crystallogr 66, 213–221 (2010).

34. S. Cogliati, J. A. Enriquez, L. Scorrano, Mitochondrial Cristae: Where Beauty Meets Functionality. Trends Biochem Sci 41, 261–273 (2016).

35. J. Habersetzer et al., ATP synthase oligomerization: from the enzyme models to the mitochondrial morphology. Int J Biochem Cell Biol 45, 99–105 (2013).

36. R. Guo, S. Zong, M. Wu, J. Gu, M. Yang, Architecture of Human Mitochondrial Respiratory Megacomplex I2III2IV2. Cell 170, 1247–1257 e1212 (2017).

37. E. Kovacs-Bogdan et al., Reconstitution of the mitochondrial calcium uniporter in yeast. Proc Natl Acad Sci U S A 111, 8985–8990 (2014).

38. M. F. Tsai et al., Dual functions of a small regulatory subunit in the mitochondrial calcium uniporter complex. Elife 5, (2016).

39. K. J. Kamer, W. Jiang, V. K. Kaushik, V. K. Mootha, Z. Grabarek, Crystal structure of MICU2 and comparison with MICU1 reveal insights into the uniporter gating mechanism. Proc Natl Acad Sci U S A 116, 3546–3555 (2019).

40. Y. Xing et al., Dimerization of MICU Proteins Controls Ca(2+) Influx through the Mitochondrial Ca(2+) Uniporter. Cell Rep 26, 1203–1212 e1204 (2019).

41. K. J. Kamer, Z. Grabarek, V. K. Mootha, High-affinity cooperative Ca(2+) binding by MICU1-MICU2 serves as an on-off switch for the uniporter. Embo Rep 18, 1397–1411 (2017).

42. G. Csordas et al., MICU1 Controls Both the Threshold and Cooperative Activation of the Mitochondrial Ca2+ Uniporter. Cell Metab 17, 976–987 (2013).

43. K. J. Kamer, V. K. Mootha, MICU1 and MICU2 play nonredundant roles in the regulation of the mitochondrial calcium uniporter. EMBO Rep 15, 299–307 (2014).

44. L. Wang et al., Structural and mechanistic insights into MICU1 regulation of mitochondrial calcium uptake. EMBO J 33, 594–604 (2014).

45. Y. Lee et al., Structure and function of the N-terminal domain of the human mitochondrial calcium uniporter. Embo Rep 16, 1318–1333 (2015).

46. S. K. Lee et al., Structural Insights into Mitochondrial Calcium Uniporter Regulation by Divalent Cations. Cell Chem Biol 23, 1157–1169 (2016).

47. T. Yamamoto et al., Analysis of the structure and function of EMRE in a yeast expression system. Biochim Biophys Acta 1857, 831–839 (2016).

48. T. Yamamoto et al., Functional analysis of coiled-coil domains of MCU in mitochondrial calcium uptake. Biochim Biophys Acta Bioenerg, 148061 (2019).

49. G. Bhosale et al., Pathological consequences of MICU1 mutations on mitochondrial calcium signalling and bioenergetics. Biochim Biophys Acta Mol Cell Res 1864, 1009–1017 (2017).

50. K. Mallilankaraman et al., MICU1 Is an Essential Gatekeeper for MCU-Mediated Mitochondrial Ca2+ Uptake that Regulates Cell Survival. Cell 151, 630–644 (2012).

51. K. J. Kamer, Z. Grabarek, V. K. Mootha, High-affinity cooperative Ca2+ binding by MICU1-MICU2 serves as an on-off switch for the uniporter. Embo Rep 18, 1397–1411 (2017).

52. A. Goehring et al., Screening and large-scale expression of membrane proteins in mammalian cells for structural studies. Nat Protoc 9, 2574–2585 (2014).

53. S. Q. Zheng et al., MotionCor2: anisotropic correction of beam-induced motion for improved cryo-electron microscopy. Nat Methods 14, 331–332 (2017).

54. J. A. Mindell, N. Grigorieff, Accurate determination of local defocus and specimen tilt in electron microscopy. J Struct Biol 142, 334–347 (2003).

55. K. Zhang, Gctf: Real-time CTF determination and correction. J Struct Biol 193, 1–12 (2016).

56. E. F. Pettersen et al., UCSF Chimera--a visualization system for exploratory research and analysis. J Comput Chem 25, 1605–1612 (2004).

57. S. H. Scheres, S. Chen, Prevention of overfitting in cryo-EM structure determination. Nat Methods 9, 853–854 (2012).

58. P. B. Rosenthal, R. Henderson, Optimal determination of particle orientation, absolute hand, and contrast loss in single-particle electron cryomicroscopy. J Mol Biol 333, 721–745 (2003).

59. A. Kucukelbir, F. J. Sigworth, H. D. Tagare, Quantifying the local resolution of cryo-EM density maps. Nat Methods 11, 63–65 (2014).

60. I. W. Davis et al., MolProbity: all-atom contacts and structure validation for proteins and nucleic acids. Nucleic Acids Res 35, W375–383 (2007).

61. O. S. Smart, J. G. Neduvelil, X. Wang, B. A. Wallace, M. S. Sansom, HOLE: a program for the analysis of the pore dimensions of ion channel structural models. J Mol Graph 14, 354-360, 376 (1996).

62. N. Alexander, N. Woetzel, J. Meiler, bcl::Cluster : A method for clustering biological molecules coupled with visualization in the Pymol Molecular Graphics System. IEEE Int Conf Comput Adv Bio Med Sci 2011, 13–18 (2011).

63. F. A. Ran et al., Genome engineering using the CRISPR-Cas9 system. Nat Protoc 8, 2281–2308 (2013).

